# Population structure and reproduction of the alvinocaridid shrimp *Rimicaris exoculata* on the Mid-Atlantic Ridge: variations between habitats and vent fields

**DOI:** 10.1101/2021.06.27.450066

**Authors:** Iván Hernández-Ávila, Marie-Anne Cambon-Bonavita, Jozée Sarrazin, Florence Pradillon

**Affiliations:** Ifremer, REM/EEP, Laboratoire Environnement Profond, F 29280 Plouzané, France; Univ Brest, Ifremer, CNRS, Laboratoire de Microbiologie des Environnements Extrêmes, F 29280 Plouzané, France; Facultad de Ciencias Naturales, Universidad Autónoma del Carmen, Ciudad del Carmen, Mexico

**Keywords:** Life cycle, population structure, reproduction, *Rimicaris exoculata*, habitat variability, hydrothermal vents

## Abstract

*Rimicaris exoculata* is a dominant species of deep Mid-Atlantic Ridge (MAR) vent fields and inhabits areas close to vent emissions at depths below 2000 m. Its high abundance and strong genetic connectivity along the MAR point at a remarkable ability to produce dispersing larval stages. However, the reproduction of this species long remained enigmatic because brooding females were rarely observed. Here, we describe the population structure and reproduction of *R. exoculata* at the Snake Pit and TAG vent fields (3600m depth) for the months of January-February. We observed major differences in population structure between habitats within a single vent field: females widely dominate the large swarms near active venting while inactive peripheries are inhabited by large males. Low temperature diffusion zones are mainly colonized by small juveniles of *R. chacei* instead of *R. exoculata*. Size structure of populations from dense active areas is polymodal at both fields, suggesting discontinuous recruitment. Male and female sizes did not vary across habitats and vent fields, with sexually mature female being slightly larger than males. In contrast to previous studies, hundreds of ovigerous females were observed at both vent fields, suggesting seasonal reproduction. Proportion of ovigerous females among sexually mature females were similar between vent fields (36.7 %). However, reproductive output was lower at TAG, where ovigerous females had smaller size-specific fecundity and egg size, and more aborted broods. Broods were colonized by the dirivultid copepod *Stygiopontius pectinatus* at both vent fields, apparently without deleterious effect on egg development. In the light of the observed variability in *R. exoculata* population structure, we propose a hypothetical scenario depicting its mating system and brooding behavior, and discuss more generally intraspecific interactions during its benthic life stages.

## 1. Introduction

The fragmentary and ephemeral nature of deep-sea hydrothermal vent ecosystems is challenging for animal populations connectivity and resilience. Visually dominant species are usually endemic to these ecosystems, and large scale distribution and connectivity between populations were observed for some of them (Thaler et al. 2011, Teixeira et al. 2012, Beedessee et al. 2013). Advances in understanding mechanisms and processes related to reproduction, dispersal, recruitment and structure of vent populations are still hampered by the difficulties to upscale our observations temporally and/or spatially due to the cost and technical challenges of deep-sea studies. Remaining gaps in our understanding of vent species life histories may hide complex population structures that have not yet been acknowledged, even in some well-studied iconic species such as vent shrimps. Indeed, a recent study in another vent crustacean, the anomuran crab *Kiwa tyleri*, revealed complex life stages distribution affecting local population structures and reflecting life history of the species (Marsh et al. 2015).

The alvinocarid shrimp *Rimicaris exoculata* is a visually dominant species at hydrothermal vents of the Mid-Atlantic Ridge (MAR), especially in fields located at depths below 2000 m. This species lives close to the vent emission in dense aggregations of thousands of individuals per square meter (Desbruyères et al. 2000, Copley et al. 2007, Gebruk et al. 2010) and many aspects of its biology have been studied since the discovery of MAR hydrothermal vents in 1985 (Zbinden & Cambon-Bonavita 2020). The relationship of this shrimp with the dense and diverse symbiotic microbial communities hosted in its cephalothoracic cavity and gut have been extensively studied, demonstrating a clear trophic dependency of the shrimp on its symbionts, and pointing at tight regulation along its life cycle (Corbari et al. 2008, Ponsard et al. 2013, Jan et al. 2014, Le Bloa et al. 2020). However, the information available about reproduction and population biology is still scattered and sometimes contradictory. In terms of population structure, major differences in shrimp densities have been found at the Broken Spur vent field depending on the levels of hydrothermal activity, including the decrease of adult densities in vent areas not directly exposed to vent emission (Copley et al. 1997). Habitats at the base of vent edifices were suggested to serve as nurseries for juvenile recruitment due to their high densities in these areas (Komai and Segonzac 2008). However, it remained unclear whether variations in adult density between habitats also involved variations in population structures. Samples collected at various MAR vent fields by Shank et al. (1998) showed a female-biased sex ratio, but the association of sex ratio with a particular habitat or temporal variation was not tested by additional sampling. In addition, patches of juveniles have been reported in adult aggregations (Shank et al. 1998, Copley et al. 2007) but their distribution and proportions in the populations remain undetermined. Other aspects of population structure, such as size structure, are based on single sample, pooled samples or preliminary analyses (Gebruk et al. 1997, Vereshchaka 1997).

The reproduction of *R. exoculata* also has many intriguing gaps. Although oocyte size frequency suggests a lack of seasonality in the reproduction (Ramirez-Llodra et al. 2000, Copley et al. 2007), very few brooding specimens have been collected since the species description (Williams & Rona 1986, Ramirez-Llodra et al. 2000, Gebruk 2010, Guri et al. 2012). The lack of brooding females also contrasts with the large densities and strong genetic connectivity observed in MAR vent fields (Teixeira et al. 2012), which must be supported by a large larval pool. Egg size has been estimated in only two studies (Williams & Rona 1986, Ramirez-Llodra et al. 2000) and this size estimation was used to infer that these shrimp should have planktotrophic larvae with long planktonic duration. However, recent morphological evidence on hatching larval stages lacking functional mouth structures led to the revision of the former hypothesis and proposition that these larvae are lecithotrophic early in their dispersive planktonic phase (Hernandez-Avila et al. 2015). In addition, the realized fecundity (988 eggs per brood) was estimated on only one specimen due the lack of samples (Ramirez-Llodra et al. 2000).

Very few data concerning the distribution of *R. exoculata* populations at different spatial scales or their interactions in vent ecosystems are available in the literature. At vents, several studies have shown that variations of physical and chemical conditions of vent emission shape the structure of communities dominated by bathymodiolin mussels or siboglinid tube worms at various spatial and temporal scales (Sarrazin et al. 1997, 2015, Cuvelier et al. 2009, 2014). Variations in community structure are observed between edifices at the field-scale (Desbruyères et al. 2000, 2001, Sarrazin et al. 2020) as well as between habitats within the same edifice (Cuvelier et al. 2009, Sarrazin et al. 2015) highlighting the high level of complexity of these ecosystems.

In addition to variations at community level, vent fauna also show variations in their population structures. These variations are linked to differences in habitats occupied by the different life stages for motile organisms (Shank et al. 1998, Marsh et al. 2015), changes in population structure between vents (Nye et al. 2013) or habitats (Copley & Young 2006, Marsh et al. 2015) and various scales of temporal variations (Copley et al. 1997, 1999, 2007, Gebruk et al. 2010, Cuvelier et al. 2011). Although less studied, there is evidence that populations of dominant species exhibit spatial and temporal variations at small scales. For instance, Copley et al. (1999) proposed that short-term changes in *R. exoculata* population density close to vent emission could be associated with tidal variation, as observed in other mobile vent species (Lelièvre et al. 2017). Variations in population structure most likely reflect the complex life cycles of vent species with different reproductive strategies, variations in physiological tolerance and resource use by the different developmental stages, and their interactions with environmental variations.

As one of the dominant species at MAR vent fields below 2000 m depth, *R. exoculata* plays a major role at ecosystem level, and its population biology could have important implications for the resilience, structure and biomass of these vent communities (Desbruyères et al. 2000, 2001). In January-February 2014, large numbers of ovigerous females were observed at two vent fields of the MAR, TAG and Snake Pit, during the BICOSE cruise. In addition, striking variations in the densities of adult and juvenile shrimps locally led us to hypothesize local variations in population structure. In this study, we tested such hypothesis and analyzed both population structure and reproductive features of ovigerous *R. exoculata* females in a series of samples collected in visually distinct shrimp assemblages. We untangled different levels of spatial variation in reproductive parameters and population structure and used this information to propose a scenario depicting interactions of shrimps with their conspecifics and their environment that would explain the observed distribution patterns, and provide clues on some aspects of their reproductive behavior and life cycle.

## 2. Material and Methods

### 2.1. Sampling

*Rimicaris exoculata* were collected at the Snake Pit (SP, 23°22.1’N 44°57.1’W, 3470 m depth) and TAG (26°08.2’N, 44°49.5’W, 3620 m depth) vent fields on the MAR (Fig. 1a) during the BICOSE cruise (DOI: 10.17600/14000100) from January 10^th^ to February 11^th^, 2014. The two vent fields are approximately 310 km away, and separated by the Kane Fracture Zone. At Snake Pit, six samples were collected in dense shrimp aggregations on the walls of active chimneys of the Beehive site, close to fluid emissions (therein Active Emission Habitat/AEH) (Table 1, Suppl. Fig. 1). At TAG, three samples were collected in the AEH of Active Mound (Fig. 1b, Suppl. Fig. 1) and two samples were collected in dense aggregations of small alvinocaridid juveniles settled in flat areas of diffuse flow (Diffuse Emission Habitat/DEH) herein termed “nurseries” (Fig. 1d, Suppl. Fig. 2). Additionally, three other samples were collected at the base of the TAG mound, where no active emission was visible (Inactive Emission Habitat/IEH) and with adult shrimps scattered over large areas (Fig. 1c, Suppl. Fig. 2). These different types of habitats occurred within meters or tens of meters from each other.

**Figure 1.**
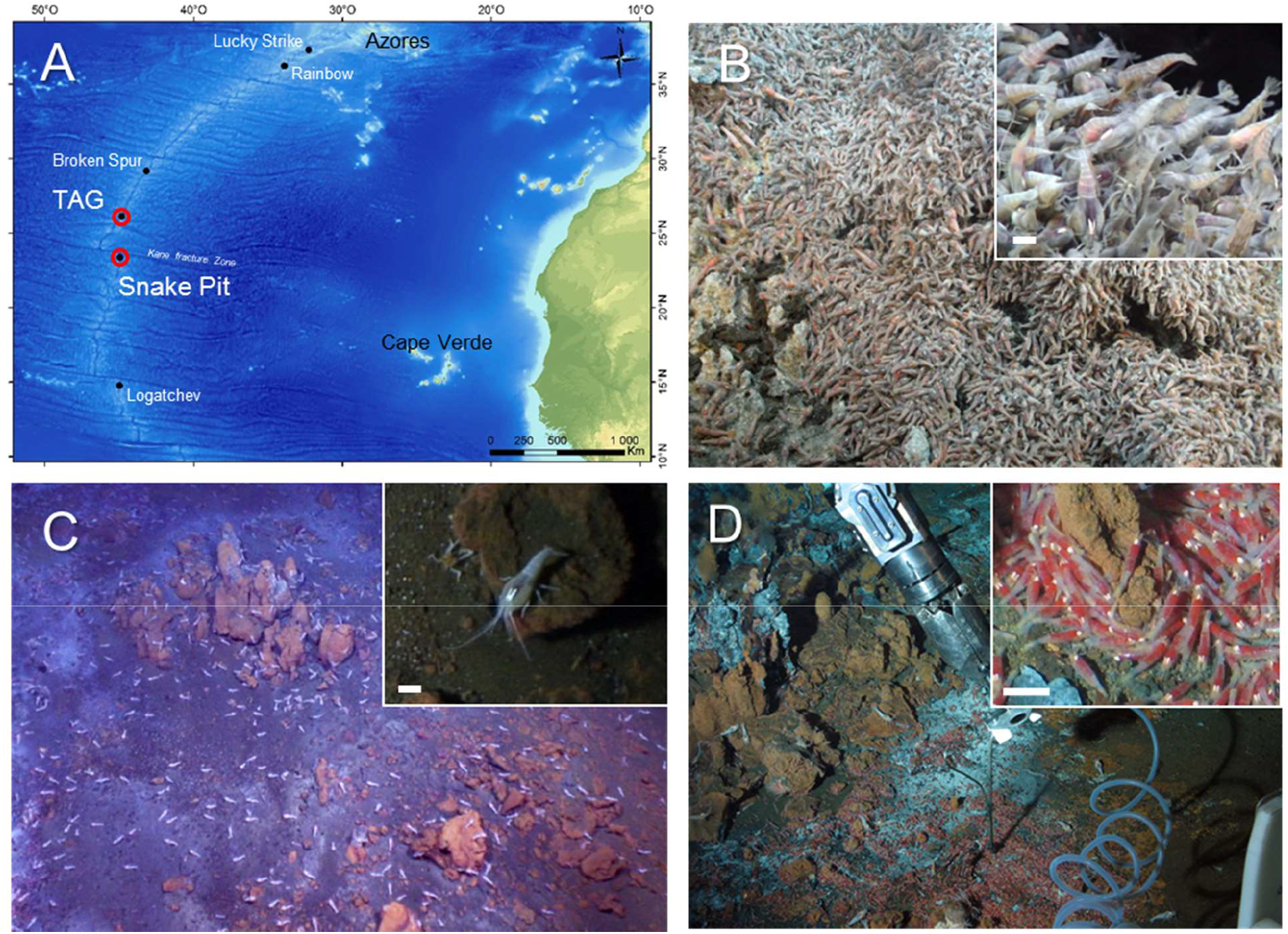
Sampled vent fields and habitats. A) North Atlantic regional map with TAG and Snake Pit vent fields on the Mid Atlantic Ridge (Amante, C. and B.W. Eakins, 2009. doi:10.7289/V5C8276M). B) *R. exoculata* swarms in the active emission habitat (AEH). C) Inactive emission habitat (IEH) with scattered *R. exoculata* adults (white spots). D) Diffuse emission habitat (DEH), showing red aggregation of small juveniles. B-D : Pictures from the TAG vent field, Ifemer/ROV Victor 6000/BICOSE2014. Scale bars in close-up views in insets: 1 cm.

**Table 1.**
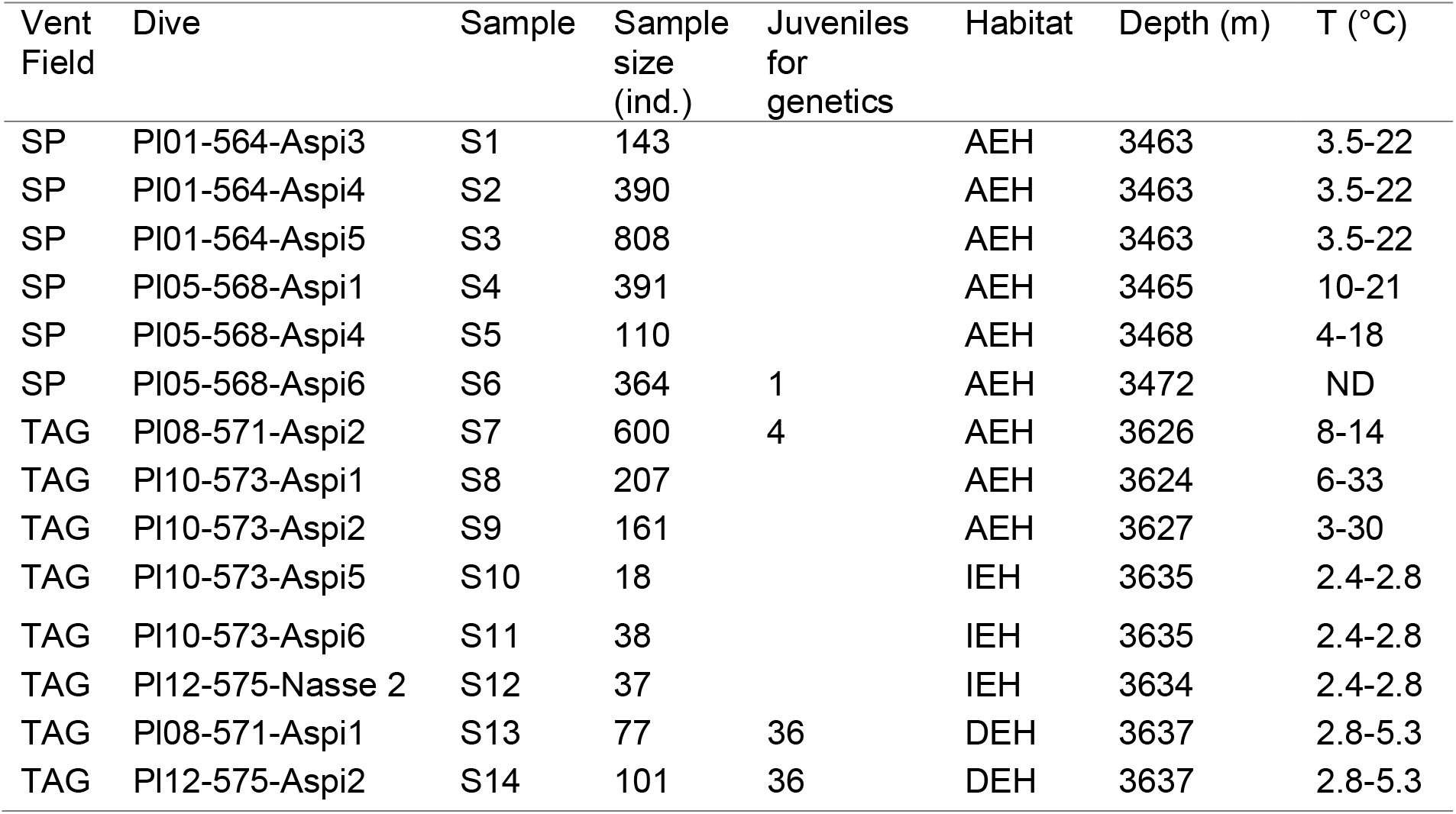
Shrimp sample details, including temperature ranges measured within centimeters from the faunal sampling point when available. ND: no data available.

Shrimps were collected with the suction sampler of the Remotely Operated Vehicle (ROV) Victor6000. In AEH and DEH, the tip of the sampler was pointed as close as possible to the individuals and maintained immobile during sampling to avoid as much as possible disturbing the aggregation. The suction was activated for a few seconds in order to collect individuals from a restricted area. Due to the low shrimp density in IEH, samples were obtained from larger areas covering a few m^2^. In addition, 37 specimens caught with a shrimp-trap deployed in IEH were included in the analyses. In total 3 445 specimens of *R. exoculata* were examined (Table 1).

Temperature measurements were conducted along with shrimp collections. Records were obtained either from discrete measurements with the submersible temperature probe prior to sampling, or from time-series measurements with autonomous temperature probes (WHOI-MISO low temp-ONSET®) deployed within the shrimp aggregation for a few days prior to sampling.

### 2.2. Identification and measurements

Specimens were initially identified and classified as juveniles, subadults or adults, in accordance with Komai & Segonzac (2008). The identification of the smaller juveniles from DEH was further assessed by DNA analyses because their morphology did not fit completely the description of early juvenile stages of *R. exoculata* available at the time of this study (Komai & Segonzac, 2008). Thirty-six juveniles from each DEH sample and 5 juveniles from AEH samples were used for molecular identifications (Table 1 & Suppl. Table 1).

Sex was identified in adults by the occurrence of the “appendix masculina” on the second pleopod in males, and the shape of the endopod of the first pleopod (Komai & Segonzac, 2008). Since these sexual characters appear during the transition from subadult to adult stage, sex of juvenile and subadult specimens could not be determined. Brooding females are characterized by the presence of embryos held between their pleopods under the abdomen, and by modifications of their pleopods (addition of setae to maintain the brood). Hatched females (females just after larval release but prior to molt) were identified by their modified pleopods. Brooding and hatched females are referred to as ovigerous females (following Nye et al. 2013). Young small adults and subadults resemble females (*i.e.* lack of appendix masculina) and also in some cases lack gonadal tissue that could be used for sex determination. In order to estimate the minimal size for confident determination of sex, a subsample of adult and subadult shrimps in small size classes (7 to 15 mm carapace lengths) were dissected to verify macroscopic evidence of gonadal tissue and estimate the onset size of sexual differentiation (OSD, see section 2.4, equivalent to the “minimum sexable size” of Anger & Moreira, 1998).

Carapace length (CL) was measured from the posterior margin of the ocular shield to the mid-posterior margin of the carapace in adults and subadults (Fig. 2a), with a precision of 0.1 mm.

**Figure 2.**
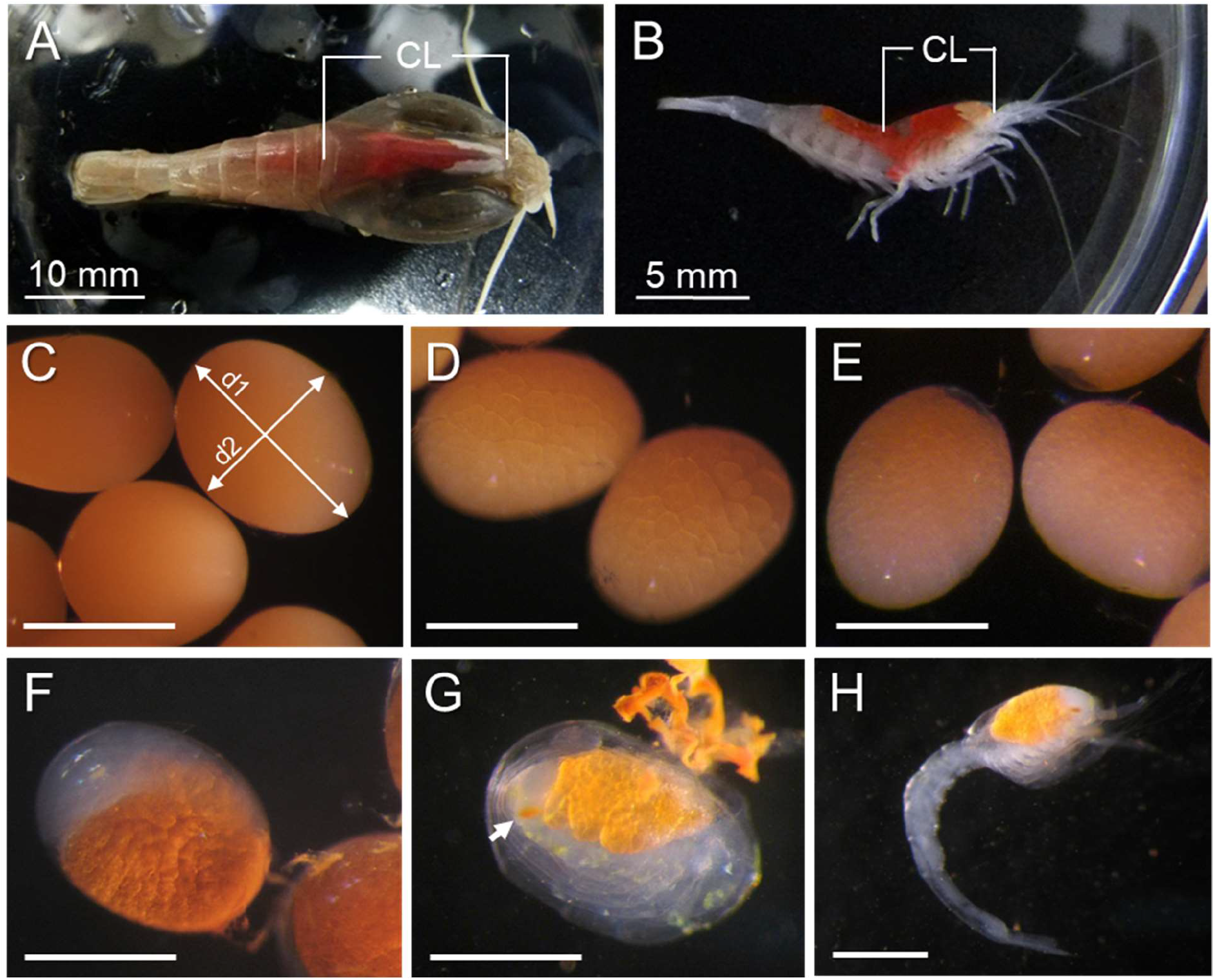
Size measurements of different life stages of *Rimicaris exoculata*. A) Adults and subadults. B) Juveniles. C-H) Embryos and hatched larvae. Early developmental stage: C) Fertilized egg (with position of measurements for maximum (d1) and minimum (d2) diameter), D) Blastula stage; Mid developmental stage: E) Early gastrulation, F) Nauplius stage; Late developmental stage: G) Pre-hatch embryo (arrow indicates eye spot). H) Hatching Zoea larva. CL: carapace length. Scale bars C-H: 500 μm.

In juvenile stages, CL was measured from the anterior tip of the rostrum to the posterior margin of the carapace (Fig. 2b). Morphological changes between juvenile and adult stages (rostrum reduction and development of the ocular shield) may introduce a bias in our measurements but this was small compared to the total length of the carapace, and had little impact on size frequency distributions and size comparisons.

Embryos were removed from the abdomen of brooding females, counted and staged. We classified embryonic developmental stages into three categories, similar to those defined by Nye et al (2013) for *R. hybisae*. Early stage embryos encompass freshly laid fertilized eggs without cellular division, and dividing eggs until the blastula stage (Figs. 2C, D). Mid stage starts with gastrulation when a clear region differentiates at one pole of the embryo and extends to the end of the naupliar development (Figs. 2E, F). Late stage includes post-naupliar development, when eye pigmentation becomes visible, abdomen appears clearly separated from the rest of the body with yolk in the cephalothorax, and appendages are fully lengthened and appear curled around the cephalothorax (Fig. 2G). For each brood, ten embryos were randomly selected and both their maximum and minimum diameters were measured at a precision of 0.03 mm under a stereomicroscope with a graduated ocular. The volume of embryos was estimated according to Oh & Hartnoll (2004), considering a spheroid volume as v= (4/3) Π r_1_r_2_^2^, where r_1_ and r_2_ are half of the maximum and minimum axis, respectively. This estimation has a precision of 1.6 x 10^−5^ mm^3^.

During examination of female broods, we found dirivultid copepods between the embryos or attached at the base of the pleopods. They were identified to the species level through barcoding (Suppl. Table 1).

### 2.3. Genetic identifications

DNA was extracted from shrimp juveniles and copepods using the CTAB method (Doyle 1990) on muscle tissue or on whole specimens for copepods. A section of the cytochrome oxidase I gene (COI) was amplified in a 50 μL solution of 1X reaction buffer, 2 mM MgCl_2_, 0.25 mM dNTP, 1.2 units of Taq polymerase and 0.6 mM of each primer (LCOI1490 and HCOI2198, Folmer et al. 1994). Amplifications were performed as following: initial denaturation (5 min at 95°C), 40 cycles including denaturation (1 min at 94°C), annealing (1 min at 52 °C) and extension (2 min at 72 °C), followed by a final extension of 7 min at 72°C. All PCR amplifications were conducted on a GeneAmp PCR system 9700 (Applied Biosystems). PCR products were purified and sequenced by Macrogen, Inc. (Netherlands) using the amplification primers. We aligned our sequences using MUSCLE (Edgar 2004), along with a set of alvinocaridid and other shrimps sequences. For our dirivultid copepod sequence, a subset of the sequences obtained by Gollner et al. (2011) was used for comparison. Neighbor-joining trees were constructed using Geneious R8 software (Kearse et al. 2012) using a HKY evolutionary model of nucleotide substitution. Robustness of the inferred trees was evaluated using bootstrap method with 1000 replicates.

### 2.4. Data analysis

Onset of sexual differentiation (OSD) was estimated using a similar procedure as what was used for determination of the size of sexual maturity in Crustacea (Wenner et al. 1974). Proportions of specimens with gonad tissue were estimated for size classes between 7-15 mm CL (larger specimens with clear sex differentiation were not included). The proportion of specimens with gonad differentiation were plotted against size classes and fitted to the logistic equation:

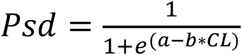

where *Psd* is the proportion of shrimps with gonad differentiation, CL is the carapace length, and *a* and *b* are constants. The size at which 50 % of the individuals exhibit sexual differentiation denoted by gonad development was considered as an estimation of OSD (Suppl. Fig. 3). Subadults were subsequently identified as individuals resembling adults with sizes < OSD.

Similarly, the size of effective sexual maturity (ESM) for females was estimated from the proportion of ovigerous females per size class, corrected by the maximum proportion of ovigerous females (King 2007). We applied the previous logistic equation, substituting *Psd* by the corrected proportion of ovigerous females (*Povf*) on the equation. The ESM was estimated by the size at which *Povf* is 50% (Suppl. Fig. 4).

For each sample, juvenile ratio, subadult ratio and sex ratio (F:M) were estimated. Deviation from a sex ratio of 1:1 was tested with a χ^2^ test, using Yates correction in samples with few specimens (n < 30). In order to determine the variations in sex ratio within each habitat at each vent field, we performed a heterogeneity χ^2^ test (Zar 2010). Variations in the proportions of females with size > ESM and the proportions of ovigerous females between vents were tested using χ^2^ test.

Size class structures were analyzed for each sample, estimating kurtosis and skewness for size class aggregation and deviation from the mean, respectively (Zar 2010). Although histograms were elaborated denoting juveniles, subadults, males and females, size structure comparisons were performed including all specimens. Normality tests were performed for each sample using the Shapiro-Wilk test (Zar 2010). Discrete size cohorts in the samples were identified using the statistical package mixdist (MacDonald 2003) running also in R™. The goodness of fit of the identified size cohorts was verified using χ^2^ test. The analyses were performed for each sample collected in the AEH and compared to verify the consistency of identified cohorts. Identification of cohorts in other habitats were not performed due to insufficient sample size.

For AEH samples, differences in body sizes associated with sex and vent fields were tested using multifactorial analysis of variance (ANOVA), after log10 transformation. For this analysis, samples were nested at the factor vent field, representing the variation in body size between samples collected within the same vent field. Similarly, an ANOVA test was performed in order to examine differences in male body sizes between habitats (AEH vs DEH) at the TAG vent field. In this case, samples were nested at the factor habitat which represents the small-scale variation. Data normality and homoscedasticity were tested using χ^2^ for frequency distribution and Levane test respectively (Underwood 1997, McGuinness 2002).

In crustaceans, brood size increases proportionally with female size (Oh & Hartnoll 2004). In order to compare reproductive outputs at the two vent fields, we estimated the relation between realized fecundity (number of embryos per brood) and female size (CL) using regression analyses, after loge transformation. The difference in size-specific fecundity between vent fields was tested using a t-test analysis. Variations in embryo size associated with parental female, embryo stage and vent fields were analyzed with a multifactorial ANOVA test. For this analysis the factor parental female was considered as nested to the combination of vent field and embryo stage.

## 3. Results

### 3.1. *Rimicaris* shrimp populations at TAG and Snake Pit in January 2014

Overall, 3445 individuals from 14 samples collected at both vent fields were examined. Among these, we determined developmental stage and sex for 3388 individuals and measured 3379 individuals (missing data are due to body damages preventing accurate measurements or sex/stage determination). The global dataset includes 1925 females (56.8%), 292 males (8.6%), 882 subadults (26.1%) and 289 juveniles (8.5%). Global sex-ratio clearly deviates from 1:1 (χ^2^, df= 1, p< 0.01). Of the 1925 females, 136 (7.1%) are either brooding eggs (125), or have recently hatched larvae (11).

Smaller specimens are early juveniles (CL = 4.4 mm), whereas the largest one is a female with 24.4 mm CL (but being an outlier specimen as other largest females reach ~20.6 mm CL). Size-range of the juveniles varies between 4.4 and 10.3 mm, and overlaps subadult size range (7-9.9 mm CL) (Suppl. Table 2). Subadult maximal size was determined by our estimation of the onset size of sexual differentiation (OSD = 10 mm, Suppl. Fig. 3). The OSD value is also consistent with the size of the smaller adult male found in our samples (CL = 9.9 mm). Although the size of some juveniles exceeds the OSD size, they are morphologically distinct from subadults: their rostrum is not completely reduced, and their carapace not fully inflated.

Overall, size ranges of males and females are similar with respective CL ranges = 9.9-19.1 mm and 10 -24.4 mm (Suppl. Table 2). However, the average size of males was higher than the average size of females, with 15.1 mm CL and 12.5 mm CL respectively (t-test= 20.71, p< 0.001). Most ovigerous females exhibit large sizes, with CL ranging from 12 mm to 20.6 mm (average size: 16.5 mm CL). We estimate the size at effective sexual maturity (ESM) at 15.1 mm CL for females (Suppl. Fig. 4). Hereafter, females with size ≥ 15.1 mm CL are called sexually mature females.

### 3.2. Variation in population structure across habitats and vent fields

#### 3.2.1. Different types of habitats

Visually, striking differences characterized shrimp assemblages when we collected our 14 samples. Most samples (9) were collected among dense aggregations of actively swimming shrimps gathering around vigorous fluid emissions (Fig. 1-B, Suppl. Fig. 1). In these aggregations, we recorded steep temporal variations in temperature with maximum varying from 14 to 33°C (Table 1). Resting shrimps scattered on inactive sulfide substratum at the periphery of dense aggregations were collected 3 times at TAG (Fig. 1-C, Suppl. Figs. 2 A-C). In this habitat, no fluid exit was visible and temperature was low and stable, with a maximum of 2.8°C (Table 1), whereas ambient seawater temperature was 2.6°C. At TAG, aggregations of very small individuals characterized by their bright red color were sampled around diffusions of translucent fluids exiting from very small chimneys or cracks (Fig. 1-D, Suppl. Figs. 2 E, F). Temperatures among those young individuals varied from 2.8°C to 5.3°C (Table 1). We thus defined 3 types of aggregations characterized by different temperature regimes, with visually distinct shrimp assemblages in terms of density, size and behavior. These are called Active Emission Habitat (AEH), Inactive Emission Habitat (IEH) and Diffuse Emission Habitat (DEH).

#### 3.2.2. Variations in population structure between habitats (TAG)

At TAG, we observe striking differences in terms of population structure, size-frequency distribution, and reproductive features among the three habitats. In AEH, with 71% of females and 8,5% of males, sex-ratio is clearly biased for females (8.4:1). Although sex ratios are significantly different among AEH samples from TAG (χ ^2^*het*= 50.05, df= 2, p< 0.001, Table 2), all of them are significantly female biased (χ^2^, df= 1, p< 0.02 in all cases, Table 2).

**Table 2.** *R. exoculata* specimens and sex ratios at different vent fields and habitats. J: juveniles, <OSD (onset of sexual differentiation): subadults, F: females (non-brooding); OF: ovigerous females; M: males.

| Active Emission<br>Habitat (AEH) |  |  |  |  |  |  |  |  |  |  |
| --- | --- | --- | --- | --- | --- | --- | --- | --- | --- | --- |
| Vent field | Sample | J | <OSD | F | OF | M | n | F : M | $\chi^2$ | p |
| Snake Pit | S01 | 1 | 8 | 109 | 1 | 20 | 139 | 5.5:1 | 62.308 | <0.001 |
| Snake Pit | S02 | 14 | 175 | 181 | 2 | 9 | 381 | 20.3:1 | 157.688 | <0.001 |
| Snake Pit | S03 | 6 | 177 | 522 | 29 | 53 | 787 | 10.6:1 | 410.603 | <0.001 |
| Snake Pit | S04 | 16 | 188 | 174 | 0 | 6 | 384 | 29:1 | 156.8 | <0.001 |
| Snake Pit | S05 | 26 | 22 | 39 | 1 | 20 | 108 | 2:1 | 6.667 | 0.01 |
| Snake Pit | S06 | 21 | 143 | 158 | 22 | 19 | 363 | 9.5:1 | 130.256 | <0.001 |
| Total |  |  |  |  |  |  |  | 9.8:1 | 904.264 | <0.001 |
|  |  |  |  |  |  |  |  | Heterogeneity | 20.056 | 0.001 |
| TAG | S07 | 19 | 138 | 380 | 35 | 23 | 595 | 18:1 | 350.831 | <0.001 |
| TAG | S08 | 1 | 6 | 163 | 25 | 7 | 202 | 26.7:1 | 168.005 | <0.001 |
| TAG | S09 | 6 | 23 | 58 | 20 | 51 | 158 | 1.5:1 | 5.651 | 0.017 |
| Total |  |  |  |  |  |  |  | 8.5:1 | 472.441 | <0.001 |
|  |  |  |  |  |  |  |  | Heterogeneity | 50.046 | <0.001 |
| Inactive Emission<br>Habitat (IEH) |  |  |  |  |  |  |  |  |  |  |
| TAG | S10 | 1 | 0 | 2 | 0 | 15 | 18 | 1:7.5 | 9.941 | 0.002 |
| TAG | S11 | 0 | 2 | 2 | 1 | 33 | 38 | 1:16 | 25 | <0.001 |
| TAG | S12 | 0 | 0 | 1 | 0 | 36 | 37 | 1:35 | 33.108 | <0.001 |
| Total |  |  |  |  |  |  |  | 1:16 | 67.6 | <0.001 |
|  |  |  |  |  |  |  |  | Heterogeneity | 0.449 | 0.993 |
| Diffuse Emission<br>Habitat (DEH) |  |  |  |  |  |  |  |  |  |  |
| TAG | S13 | 77 | 0 | 0 | 0 | 0 | 77 |  |  |  |
| TAG | S14 | 101 | 0 | 0 | 0 | 0 | 101 |  |  |  |

In contrast, in IEH, 90.3% of the individuals are males, while females represent only 6.5% of the shrimps collected. Sex ratio is significantly biased for males (1:14) and in each TAG IEH samples (χ^2^, df= 1, p≤ 0.002 in all cases, Table 2).

Subadults represent 17.5% of the individuals on average in AEH (Table 2), with significant variation between samples (from 3% to 23.2% of the individuals in each sample, χ^2^*het*= 16.4, df= 2, p< 0.001), but they are rare in IEH with only two individuals collected overall.

Ovigerous females are almost exclusively observed in AEH at TAG (only one hatched female observed in IEH and none in DEH), representing 11.7 % of the females on average, with strong variations between samples (from 8.4% to 25.6%). These variations reflect both variations in the proportion of sexually mature females among all females (22.4% on average, with variations between samples: 13.3% to 64.9%), and, to a lesser extent, variations in the proportion of ovigerous females among sexually mature females (36.8 % on average, with variations between samples: 29.8% to 43.6%).

Juveniles are not abundant in AEH (less than 3% of the total population, Table 2, no significant heterogeneity between samples: χ^2^*het*= 3.64, df= 2, p= 0.602), and are rare in IEH with only one early stage juvenile collected. In contrast, DEH samples are exclusively composed of early stage juveniles. Although they were not collected, few large adult individuals (both *R. exoculata* and *R. chacei*) were observed crawling around these nurseries (*in situ* observations and Fig. 1-D).

#### 3.2.3. Variation in population structure between vent fields (AEH)

AEH Snake Pit samples exhibit similar population structure to those from TAG, with a strong dominance of females. Females, males, subadults and juveniles represent respectively 57.2%, 5.9%, 33 % and 3.9% of the overall population. Sex-ratio is female-biased (9.8:1 overall) and similar to the ratio observed in AEH at TAG. Like in TAG AEH samples, sex-ratio vary significantly among AEH samples at Snake Pit (χ ^2^*het*= 20.06, df= 5, p= 0.001, Table 2), but all are significantly female-biased (χ^2^, df= 1, p≤ 0.01 in all cases).

Overall, subadults are more abundant in Snake Pit samples than in TAG samples, representing almost a third of the population. However, their proportion varies strongly between samples (from 5.8% to 45.9% of the individuals, χ^2^*het*= 164.9, df= 5, p< 0.001). Like in TAG AEH, the proportion of juveniles in Snake Pit samples is generally low, except in one sample where they reach 24.1% of the individuals, resulting in significant heterogeneity between Snake Pit samples (χ^2^*het*= 89.32, df= 5, p< 0.001).

Although the proportion of ovigerous females among sexually mature females is similar between vent fields (36.6 % on average at Snake Pit, χ^2^= 0.003, p= 0.956), the proportion of sexually mature females among all females is significantly lower in Snake Pit (11.7 %) than in TAG (22.4 %) AEH samples (χ^2^= 34.24, p< 0.001), resulting in lower proportion of ovigerous females overall (4.4% of all females).

#### 3.2.4. Variations in size frequency distributions among habitats and vent fields

Overall, reflecting the differences in sex and stage distributions between habitats, size distributions also differed between habitats. While DEH host very small shrimps almost not represented in other habitats, mostly large individuals inhabit IEH. In AEH, shrimp sizes vary over a wide range, overlapping slightly both size ranges of shrimps in DEH and IEH (Fig. 3, Suppl. Fig. 5).

**Figure 3.**
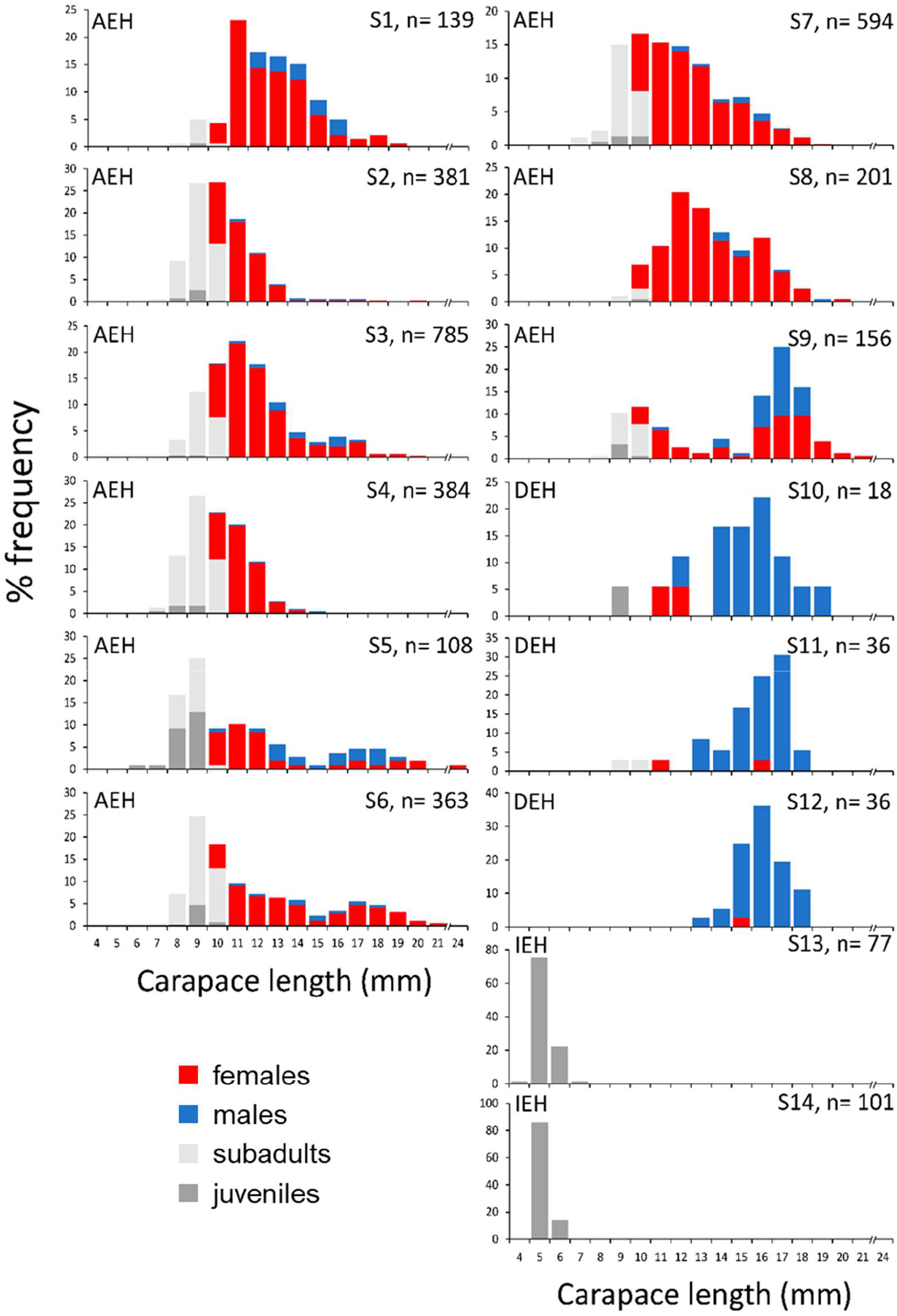
Size class structure of *R. exoculata* in different habitats of the Snake Pit (left side) and TAG (right side) vent fields. AEH active emission habitat, DEH diffuse emission habitat, IEH inactive emission habitat.

Size frequency distributions vary among samples and habitats in both vent fields, both in terms of kurtosis and skewness (Fig. 3). General trends in AEH size frequency distributions include bias towards small sizes (skewness= 0.945) and slightly leptokurtic distribution (kurtosis= 0.423). In some samples, the distribution is clearly non-unimodal. In others, deviation in skewness and kurtosis suggests a mixture of cohorts. Based on a size cohort analysis, 5 different cohorts are identified overall in AEH (Table 3). In individual samples, cohort number vary between 2 to 4 (Table 3, Suppl. Fig. 6), with some correspondence observed between samples across the two vent fields. Overall in AEH, we identify one cohort of juveniles and subadults, one cohort of subadults and small adults (<12 mm CL) and three cohorts of adults, two of which include females ≥ ESM. In both vent fields, the cohorts corresponding to juveniles, subadults and small adults represent an important proportion of the population of AEH.

**Table 3.** Identified cohorts in AEH at the Snake Pit and TAG vent fields. Mean and standard deviations are shown for each sample, proportion of each cohort are in brackets. χ^2^ denote the deviation of the sample from the cohort estimation. ns: non-significant.

| Cohort: | 1 | 2 | 3 | 4 | 5 | $\chi^2$ |
| --- | --- | --- | --- | --- | --- | --- |
| Snake Pit |  |  |  |  |  |  |
| S1 |  | 11.45±1.75<br>(0.59) | 13.24±2.03<br>(0.41) |  |  | 12.93ns |
| S2 | 8.86±0.87<br>(0.61) | 10.78±1.06<br>(0.37) |  | 16.14±1.58<br>(0.02) |  | 3.058ns |
| S3 | 9.14±0.89<br>(0.33) | 11.25±1.09<br>(0.54) |  | 15.49±1.50<br>(0.13) |  | 11.16ns |
| S4 | 8.45±0.77<br>(0.50) | 10.47±0.96<br>(0.48) | 12.92±1.18<br>(0.02) |  |  | 1.995ns |
| S5 | 8.30±0.87<br>(0.52) | 11.40±1.19<br>(0.28) |  |  | 17.28±1.80<br>(0.20) | 15.32ns |
| S6 | 8.70±0.70<br>(0.47) | 10.44±0.84<br>(0.17) | 12.81±1.03<br>(0.17) |  | 17.11±1.38<br>(0.19) | 3.247ns |
| TAG |  |  |  |  |  |  |
| S7 | 9.83±1.40<br>(0.59) |  | 13.38±1.90<br>(0.41) |  |  | 25.11* |
| S8 |  | 10.12±0.83<br>(0.15) | 12.17±1.00<br>(0.51) | 15.39±1.27<br>(0.34) |  | 3.556ns |
| S9 | 8.98±0.53<br>(0.21) | 10.50±0.62<br>(0.11) | 13.37±0.79<br>(0.06) |  | 16.73±0.99<br>(0.62) | 8.230ns |

\*Despite the deviation of the size structure from the cohort model ( $p=0.0485$ ), it was the closest model to the input data

Male and female body sizes in AEH are significantly different among samples, indicating small scale variations in sizes, within habitats at each vent field (ANOVA, p< 0.001, Suppl. Table 3). Body sizes also vary significantly with sex (ANOVA, p< 0.001, Suppl. Table 3), males being larger than females at each vent field (Suppl. Table 2). Size distribution of ovigerous females clearly departed from the rest of the females, with sizes similar to the male average size in TAG and even larger than the male average size in Snake Pit. Although males and females tend to be smaller in Snake Pit than in TAG AEH, size differences are not significant between the two vent fields for each sex (ANOVA, p= 0.083, Suppl. Table 3). In contrast, ovigerous females are slightly larger in Snake Pit than in TAG AEH, but this variation between the two vent fields was not statistically significant (ANOVA F= 1.649, p= 0.246, df_2_= 6).

Size frequency distributions of IEH samples are leptokurtic (kurtosis 0.12 to 1.99) and biased towards larger sizes (skewness −0.45 to −1.52). Males do not exhibit different body sizes between AEH and IEH, although significant differences are detected between samples of a given habitat (ANOVA, p< 0.001, Suppl. Table 4). Females also exhibit similar size ranges between both habitats, although the low number of females collected in IEH prevents statistical comparisons.

#### 3.2.5. Juvenile distribution in AEH and DEH: detection of Rimicaris chacei nurseries

Juvenile sizes at the AEH are similar between vent fields, only showing differences between samples (ANOVA F_vents_= 5.2001, p= 0.057, df_2_= 7, F_samples_= 2.643, p= 0.015, df_1_= 7, df_2_= 101). However, juveniles from DEH are much smaller (ANOVA F_vents_= 804.91, p< 0.001, df_2_= 3, F_samples_= 1.580, p= 0.195, df_1_= 3, df_2_= 199) and form a distinct cohort from that of AEH. Based on morphological features described in Komai & Segonzac (2008), we suspected that juveniles from DEH were possibly a mixture of *R. exoculata* and *R. chacei*. Molecular identification using COI barcode reveals that all sequences of these juveniles are consistent with *R. chacei* affiliation, whereas juveniles from AEH (with clear *R. exoculata* juvenile morphology according to Komai & Segonzac 2008) are consistent with *R. exoculata* affiliation (Suppl. Fig. 7).

### 3.3. Reproductive features

#### 3.3.1. Fecundity

Of the 125 brooding females found in our samples, 36 had obviously lost part of their broods, not because of hatching -since embryos were clearly not yet fully developed- but rather due to either abortion or more probably lost during sampling. These are not included in our fecundity analyses.

Fecundity varies from 304 eggs in a female from TAG with 16.2 mm CL, to 1879 eggs in a female from Snake Pit with 19.8 mm CL. The largest brooding females are observed at Snake Pit (with 1704 eggs and 20.5 mm CL for the largest), while the smallest brooding females are from TAG (with 500 and 532 eggs for the two smallest −12 mm CL- individuals). The average fecundity is overall of 833 eggs, and is higher among Snake Pit brooding females (1045 eggs, with an average CL of 17.4 mm) than among TAG ones (616 eggs, with an average CL of 15.9 mm). As expected, a positive correlation is observed between carapace length of the females and fecundity (Pearson correlation, R= 0.682, *t-test*= 8.71, p< 0.001) (Fig. 4A). We consider that more data are necessary to estimate accurate linear regression models of fecundity and compare differences between populations or with other alvinocaridid shrimps.

**Figure 4.**
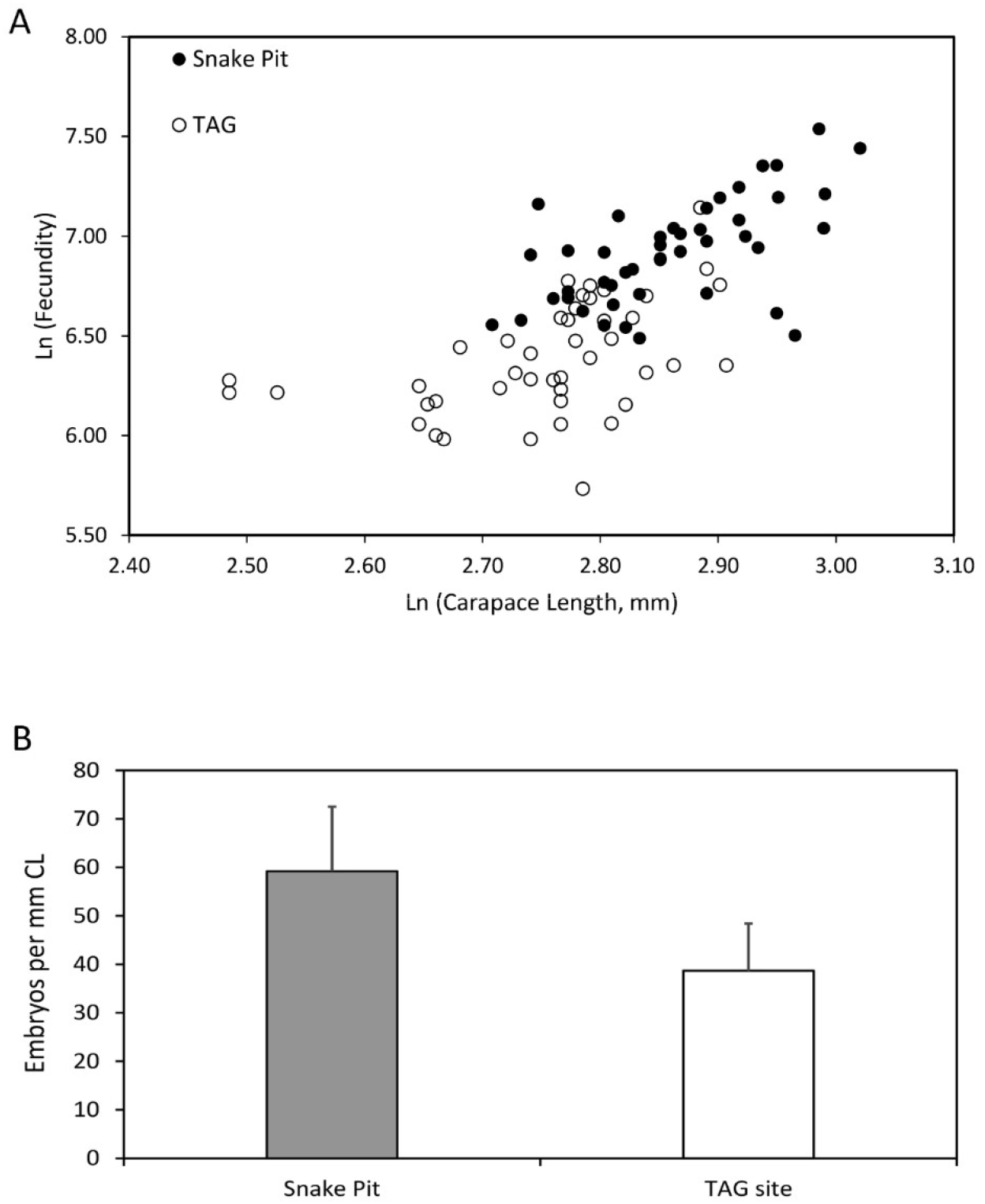
Fecundity in *R. exoculata* from TAG and Snake Pit vent fields. A. Number of embryos related with female size (carapace length). B. Size-specific fecundity, number of embryos per unit of size (mm of carapace length).

Overall size-specific fecundity ranges from 19 to 95 embryos.mm^−1^ CL. The size-specific fecundity of females does not change with the developmental stage of the brood (Early vs Mid stage, t-test= 0.98, p= 0.164, df=57; Mid vs Late stage, t-test= 0.17, p= 0.432, df=57), but is significantly lower in females from TAG (39 ± 10 embryos.mm^−1^ CL) than in females from Snake Pit (59 ± 13 embryos.mm^−1^ CL) (t-test=8.16, p<0.001, df=87) (Fig. 4B). Finally, more females with damaged broods are observed at TAG, although we could not identify the cause of loss (abortion or sampling).

#### 3.3.2. Reproductive synchrony

Within each individual brood examined, all eggs are at the same developmental stage (early, mid or late), except for occasional dead embryos or non-fertilized eggs. However, embryos at all developmental stages are observed in females from both vent fields, showing a lack of synchrony between the broods of different females. Overall, the distribution of brood stages are different between vent fields (χ^2^= 7.097 p=0.014), with some variability between samples of a given vent field (Suppl. Fig. 8). At both vent fields, a third of the females carry early stage broods, however at Snake Pit late-stage brooding and hatched females were slightly more frequent (37%) than at TAG (19%). At TAG, most of the brooding females are at the mid stage (51.9%) (Fig. 5A).

**Figure 5.**
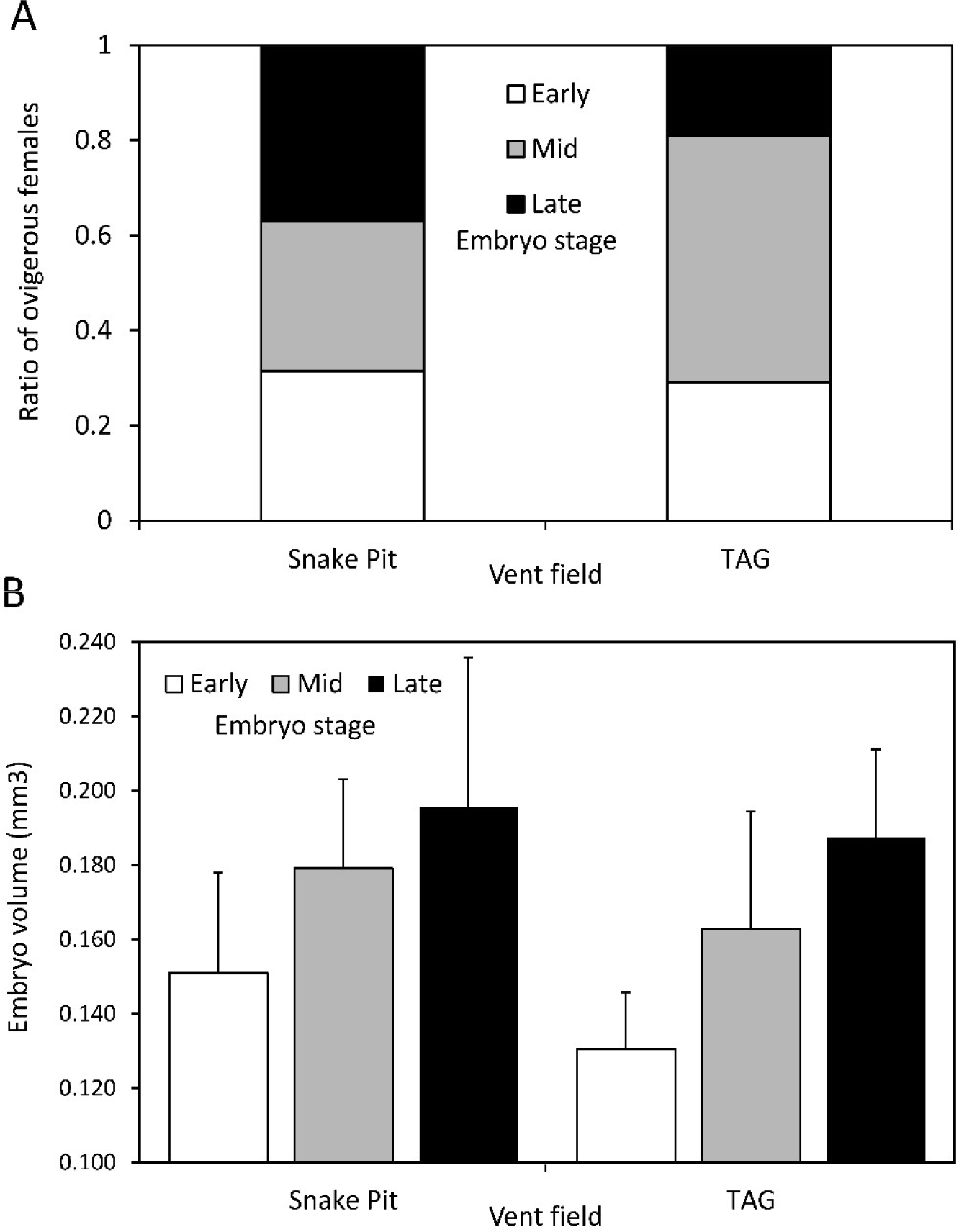
Characteristics of *R. exoculata* broods at TAG and Snake Pit vent fields. A. Proportion of broods with eggs at each developmental stage (including hatched females as having late broods). B. Sizes of individual embryos at different developmental stages.

#### 3.3.3. Egg sizes and development

The volume of the eggs within the brood of each female shows significant heterogeneity due to inter-individual variations, developmental stage of the broods, and vent fields (Supp. Table 5). Despite individual variations, a clear trend of egg volume increase with developmental stages is observed at both vent fields (Fig. 5B). The ratio between the minimum and maximum diameters of embryos decreases along their development, with minimum diameter being on average 0.87 of the maximum diameter in early stage, and 0.76 of the maximum diameter in the late stage. Eggs thus become more elongated at the end of embryonic development, which may reflect an increase of the embryo polar axis, the distribution of the structures inside the envelope and water uptake.

At each stage, embryos in TAG broods are smaller than in Snake Pit broods (Figure 5B). The volume of early stage embryos is 0.151 ± 0.027 mm^3^ at Snake Pit, and 0.131 ± 0.015 mm^3^ at TAG (*t-test*= 8.386, p< 0.001, df=242). At mid-stage, the volume of embryos increases to 0.179 ± 0.024 mm^3^ at Snake Pit and 0.163 ± 0.032 mm^3^ at TAG (*t-test*= 3.719, p< 0.001, df= 116). In the late stage, the volume of embryos reaches 0.196 ± 0.040 mm^3^ at Snake Pit and 0.187 ± 0.024 mm^3^ at TAG (*t-test*= 2.458, p= 0.007, df= 238).

#### 3.3.4. Copepod occurrence within broods

The copepods collected from the brooding females were identified as *Stygiopontius pectinatus* using molecular barcodes (Suppl. Fig. 9). These copepods appear to be truly associated with eggs because they are found deep inside the broods usually attached to the setae at the basis of the pleopods. In addition, no copepods were found on the abdomen of non-brooding females. At Snake Pit, 33 % of the brooding females are colonized with copepods, whereas only 20 % of them are infected at TAG. The proportion of infested females tends to increase between early (10%) and late (40%) stage broods (χ^2^= 6.540, p= 0.038, Suppl. Figure 10). This trend is observed at each vent field and for each sample. In infested broods, the number of copepods per brood vary between 1 and 5, and appears to be higher at Snake Pit (2 copepods per brood in average, range: 1-5 copepods) than at TAG (1.1 copepods per brood in average, range: 1-2 copepods). Similarly, the average number of copepod per brood tends to increase with brood age from 1 copepod/brood in average in early broods to 2 copepod/brood in average in late broods.

## 4. Discussion

In January-February 2014, samples of *Rimicaris exoculata* populations at TAG and Snake Pit vent fields revealed strong female-biased sex ratios (87% females among adults overall). Sex ratio biases were also striking locally, reflecting abrupt changes in population structure among habitats. Dense shrimp assemblages crawling next to high temperature fluid exits (AEH) consisted mainly of females and immature individuals, whereas shrimps observed scattered in the cold and stable periphery of active vents (IEH) were almost only adult males. At TAG, near low temperature diffusion areas (DEH), gathering of very small juveniles absent from AEH, and previously hypothesized to be early juveniles of *R. exoculata*, were indeed *R. chacei* nurseries. Ovigerous females were observed in larger proportion than ever reported so far for the species, representing about a third of the sexually mature females in AEH. Overall, these patterns were consistent across both vent fields, although a high degree of heterogeneity in population structure was observed at small spatial scales, between samples from a given habitat. Ovigerous females were more abundant at TAG, reflecting a higher proportion of sexually mature females. However, lower fecundities, smaller eggs as well as a higher proportion of aborted broods, were observed in ovigerous females from TAG, suggesting a lower individual reproductive effort.

### 4.1. Spatial variation in *R. exoculata* population structure and reproduction

Our study describes strongly biased sex ratios in *R. exoculata*, where sexes appear to segregate between different habitat types. Similar evidence has been previously reported by Shank et al. (1998) in samples collected from active chimney walls of several MAR vent fields, where females were found in much larger numbers than males. Although variations in sex ratios between the different vent fields were observed, variations within vent fields could not be assessed because unique samples were collected in each field (Shank et al. 1998). In 2007, shrimps collected in AEH at Logatchev during the Serpentine cruise also showed a female-dominated population (I. Hernández-Ávila, unpublished data). Population structure of shrimps in IEH has not been reported previously, probably because individuals scattered at the periphery of dense aggregates have been considered as remains of the main populations observed in AEH, rather than individuals preferentially occupying a specific and separated habitat. Therefore, similarly to our observations, variations in sex ratios observed in previous studies could indeed reflect small-scale local variations, rather than variations between vent fields.

Female-biased sex ratios have been reported in several vent crustaceans, often associated with different spatial patterns among sexes. For instance, Nye et al. (2013) reported that populations of *Rimicaris hybisae* from the Cayman Trough have a sex ratio in favor of females close to the vent emissions, while more dispersed populations are dominated by males at their peripheries, with some degree of local variability. In brine pools of the Gulf of Mexico, *Alvinocaris stactophila* shows overall female-biased sex ratio with males preferentially locating on the outer part of mussel beds (Copley & Young 2006). At hydrothermal vents of the East Scotia Ridge, the chirostylid crab *Kiwa tyleri* exhibits similar -but inverse- patterns: areas close to vent fluids emission are occupied by dense male-dominated aggregations, whereas females are more numerous at the periphery (Marsh et al. 2015). *A. muricola* inhabiting cold seeps off Congo also revealed a globally female-biased sex ratio, with significant local variation observed between limited numbers of samples (Ramirez-Llodra & Segonzac 2006). Other vent shrimps such as the Hippolytidae *Lebbeus virentova* from the Cayman Trough also exhibit female-biased sex ratio, but spatial variation was not reported (Nye & Copley 2014).

Immature individuals including juveniles were observed in AEH and IEH. However, both habitats were occupied by apparently different life stages, with very small juveniles in IEH, whereas AEH harbored larger juveniles and subadults. Small juveniles of IEH had inconsistent morphologies with a mixture of characters described for both early juveniles of *R. exoculata* (stage A *sensus* Komai & Segonzac 2008) and juveniles of *R. chacei*. Barcoding of these IEH juveniles revealed that they were indeed all *R. chacei*, whereas larger juveniles in AEH belonged to *R. exoculata*. This observation has been repeated in 2017 and 2018, with large aggregations of small juveniles of *R. chacei* observed around vent diffusion areas at the periphery of TAG and Snake Pit vent fields (Methou et al. 2020). A more systematic examination of morphological characters along with molecular characterization led to a redefinition of the juvenile stages of both *Rimicaris* species (Methou et al. 2020). Our nurseries here are thus specific of *R. chacei*, whereas juveniles of *R. exoculata* appear to be found only in AEH, where they grow to the subadult and adult stages. Although this is not shown in details here, juveniles corresponding to the 2 juvenile stages of *R. exoculata* described in Methou et al. (2020) occurred in our AEH samples. Variations in the proportions of juveniles and subadults in AEH may hide further structuration at very small spatial scales, perhaps corresponding to environmental gradients associated with vent fluid and seawater mixing. Additional observations are necessary to elucidate such microscale population structure within AEH.

### 4.2. Reproductive efforts of *Rimicaris* and potential environmental effects

The size for sexual differentiation (OSD) is 10 mm, about half of the maximal size of the species (not considering our single outlier). While the OSD parameter has practical value for sex separation in studies of population structure, size at first reproduction represents a key life-history parameter reflecting the life-time investment in reproduction of a species (Anger & Moreira 1998). In Alvinocaridid species, the size at first reproduction varies between 50% (*R. hybisae*, Nye et al. 2013) and 60% (*A. muricola*, Ramirez-Llodra & Segonzac 2006, *A. stactophila*, Copley & Young 2006, *M. fortunata*, Ramirez-Llodra et al. 2000) of the maximal size of the species. *R. exoculata* thus falls within the range of its family, with the smallest brooding females measuring 12 mm CL, which represents 58% of its maximal size (20.6 mm CL, not considering our outlier). However, few females between 12 mm CL and 15 mm CL (i.e. < ESM) were brooding eggs in our samples (3.5%) and likely represent ‘premature’ specimens, while much more become sexually mature (36.5%) when they reach 15.1 mm CL (ESM), which represents 73% of the species maximal size (excluding outliers). Shrimps of the family have been reported as iteroparous (Copley & Young 2006, Nye et al. 2013, Ramirez-Llodra et al. 2000), and a later onset of reproduction might limit life time investment in reproduction. On the other hand, since large females have larger broods, favoring reproduction to the largest individuals might be advantageous and help to maximize energy investment in reproduction.

Fecundity in *R. exoculata* was on average of 833 eggs/female, ranging from 304 to 1879, with brood sizes increasing with female sizes, which is common in carideans (Corey & Reid 1991). Previous counts on the rarely collected ovigerous females fall within this range. Ramirez-Llodra et al. (2000) reported a female from Snake Pit (Microsmoke cruise, November 1995) with 988 eggs (16.4 mm CL) and a female from TAG (Bravex cruise, September 1994) with 836 eggs (17.7 mm CL). *R. exoculata* appears to have fecundities similar to those reported for *R. hybisae* (maximal fecundity of 1707 eggs, Nye et al. 2013) but lower than fecundities of *R. chacei* (2510 eggs reported in a female collected in June 1994 at Lucky Strike, Ramirez-Llodra et al. 2000). Egg sizes of *R. exoculata* are also consistent with the previous report of Ramirez-Llodra et al. (2000): early stage (blastula) embryos had a volume of about 0.145 mm^3^ (volume are calculated hereafter from diameters given in reference papers, with the spheroid formula used in this paper) at TAG, within the size range we observed in this study at similar developmental stage: 0.131 mm^3^ at TAG and 0.151 mm^3^ at Snake Pit. Eggs of *R. exoculata* are larger than those of *R. hybisae* (0.08 mm^3^, Nye et al. 2013), or those of *R. chacei* (0.09 mm^3^, Ramirez-Llodra et al. 2000). They are also larger than the eggs of *A. muricola* (0.1 mm^3^, Ramirez-Llodra & Segonzac 2006) and similar in size to the eggs of *M. fortunata* (0.13 mm^3^, Ramirez-Llodra et al. 2000). *R. exoculata* thus stands among the Alvinocarididae shrimps with the largest eggs, perhaps reflecting a specific strategy of higher parental investment per egg.

We observed significant variations in reproductive outputs between *R. exoculata* from TAG and Snake Pit. With lower realized fecundities, perhaps partly due to higher post-spawning losses, and lower egg sizes, the reproductive effort of individual females at TAG indeed appears to be reduced compared to that of females from Snake Pit. The difference in fecundity could be explained by the larger body sizes of brooding females at Snake Pit, and the positive allometric variation of fecundity with female size. Similarly, differences in fecundity between *R. hybisae* females of two vent fields in the Cayman Trough were primarily attributed to the large size differences observed between shrimps of the two fields (Nye et al. 2013). However additional factors, such as food availability, environmental challenges (fluid toxicity, or temperature stress), or variations in fertilization success may also contribute to fecundity differences between both vent fields.

The smaller size of the embryos carried by females from TAG compared to those from Snake Pit suggests that the reproductive investment per egg is lower at TAG, further contributing to the lower reproductive output observed at this vent field. Both vent fields are located approximately 310 km apart nearly at the same depth, and regional factors are unlikely to provide environmental heterogeneity that could explain the different investment in reproductive efforts. We thus hypothesize that reproduction differences between the two populations are more likely related with local environmental factors associated with vent emissions. Shrimp tolerance to metallic elements and dissolved gases in vent fluids depends on detoxification processes through metallothioneins, antioxidants (Gonzalez-Rey et al. 2007) and metabolic activities of their symbiotic bacteria (Jan et al. 2014). Higher concentrations in some metallic elements in hydrothermal fluids could force the shrimps to allocate more metabolic energy to detoxification processes, at the expense of reproductive functions. For instance, TAG fluids have higher iron, cupper and manganese concentrations than those from Snake Pit (Desbruyères et al. 2000, Schmidt et al. 2008, Charlou et al. 2010), which could explain the lower reproductive output of shrimps at this vent field. However, both bioenergetics and vent processes are complex and these hypotheses must be explored with experimental approaches testing physiological tolerance of the shrimps (e.g. August et al. 2016) in relation with more detailed exploration of local vent chemistry.

*Stygiopontius pectinatus* copepods are known to inhabit the cephalothoracic cavity of *R. exoculata* (Gollner et al. 2010, Humes 1996, Ivanenko et al. 2006). They belong to the Siphonostomatoida, an order of copepods comprising many cases of parasitic lifestyles. *Stygiopontius* copepods of the Juan de Fuca Ridge graze bacteria in alvinellid tubes (Limen et al. 2008) and *S. pectinatus* are believed to feed on microbial communities developing in the cephalothoracic cavity of *R. exoculata* (Gollner at al. 2010, Humes 1996). Here, they appear to also inhabit broods, which could similarly provide shelter and food, since the egg envelops are colonized by bacteria (Methou et al. 2019). We observed a trend of increasing copepod infestation with brood aging, which also correlates with the development of microbial communities on egg envelopes, but could also simply be the result of cumulative probability of encounter with time. The infestation of Alvinocaridid broods with small meiofaunal organisms has been briefly reported for some brooding females of *A. muricola* in cold seeps of the Gulf of Guinea (Ramirez-Llodra & Segonzac 2006). Small nematodes *Chomodorita* sp. nov. were reported among the eggs of *A. muricola*, and were associated with the presence of decaying embryos, with one instance of predatory behavior (the nematode was partly inside the embryo) reported. In *R. exoculata* broods, although a more extended survey would be needed to statistically assess the impact of *S. pectinatus* copepods on egg development, our observation did not reflect a negative effect, as higher infestation rates were found at Snake Pit where broods had more numerous, larger and healthier eggs.

### 4.3. Spawning seasonality in R. exoculata?

Despite three decades of sampling at MAR hydrothermal vents, brooding females were very rarely reported in *R. exoculata* (Vereshchaka 1997, Ramirez-Llodra et al. 2000). The only exception until 2014, was the observation of brooding females on structures of the Logatchev vent field in March 2007, and the collection of a few of them (Komai et Segonzac 2008, Gebruck et al. 2010, Guri et al. 2012). In January-February 2014, large numbers of ovigerous females were observed crawling among dense shrimp aggregates on sulfide structure walls. In our samples, the overall abundance of ovigerous female was relatively low (7.1% of the sampled females) but this probably reflects the low proportion of females reaching the size of sexual maturity. In fact, while a third of the sexually mature females were ovigerous, many of the two remaining thirds exhibited bright pink well developed gonads suggesting they were nearly ready to spawn (observation not quantified, seen in our samples and on *in situ* video acquisition). This high proportion of ovigerous females among sexually mature ones, consistent across two vent fields 310 km apart, and contrasting with all previous collections along the MAR (see 1985-2014 collection compilation, Table 4) suggests temporal variations in reproductive activity of *R. exoculata* with a possible spawning season in winter.

**Table 4.**
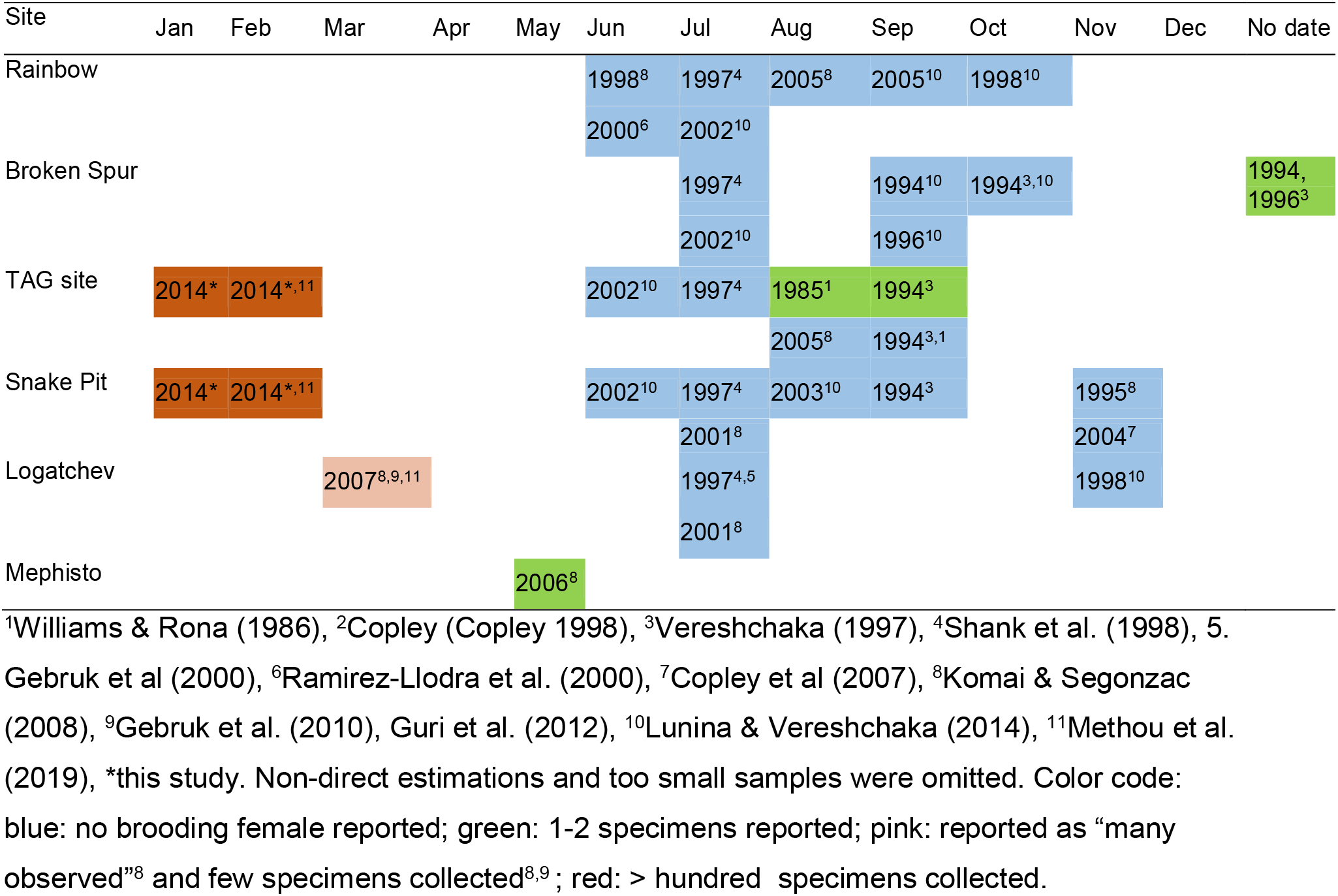
Occurrence of brooding females in *Rimicaris exoculata* samples from different cruises done on the MAR since 1985.

However, previous studies on *R. exoculata* reproduction suggested continuous reproduction based on asynchronous ovarian development observed among females collected in summer at the Rainbow vent field (Ramirez et al. 2000) and in autumn at TAG (Ramirez-Llodra et al. 2000, Copley et al. 2007). Nonetheless, regardless of vent origin, oocyte maximum size was significantly lower in summer than in autumn (Ramirez-Llodra et al. 2000). In addition, brooding females were extremely rare in both studies (3 specimens totally reported in Ramirez-Llodra et al. 2000, collected in August, September and November). This thus might suggest an extended period of oocyte growth and maturation in summer and autumn, with rare occurrences of early spawning, followed with a winter period during which large proportion of sexually mature females spawn. Such scenario is similar to what is observed in *Alvinocaris stactophila* from cold seeps in the Gulf of Mexico (Copley & Young 2006). *A. stactophila* females exhibit increasing oocyte size throughout the year, with small previtellogenic oocytes in both ovigerous and non-ovigerous females in February-March, larger vitellogenic oocytes in summer with no ovigerous female observed and females with large vitellogenic oocytes and small previtellogenic oocytes in November. In the latter, females with large vitellogenic oocytes are large non-ovigerous females (but ready to spawn), while females with small previtellogenic oocytes were either ovigerous females or small bodied non-ovigerous females (Copley & Young 2006). Such pattern fits a seasonal spawning between November and March in *A. stactophila*, and appears very similar to what is observed in *R. exoculata*. In *R. hybisae* from vents in the Cayman Trough, winter spawning was also suggested (Nye 2013, Nye et al. 2013). In winter, most females had one cohort of small previtellogenic oocytes, while a smaller number of them had a cohort of large vitellogenic oocytes, and only a few had a cohort of intermediate size oocytes. Large and medium size vitellogenic oocytes were observed only in non-ovigerous females, while small size previtellogenic oocytes were observed both in ovigerous and non-ovigerous females, suggesting that while some females had already spawned, others had mature oocytes and would probably spawn shortly (Nye 2013, Nye at al. 2013). Inter-annual comparison of the proportion of brooding females between January and February further support winter spawning in *R. hybisae*, but the effect of spatial population structure and need to expand temporal coverage was recognized (Nye 2013). Although we did not dissect ovaries in our study, we noticed that many non-ovigerous females in the large size classes had conspicuous pink-orange gonads filled with large oocytes (visible through the transparent carapace) indicative of imminent spawning, whereas ovigerous females had shrank whitish gonads, which reflects observations on ovaries of *R. hybisae* during the winter spawning period. As for *R. hybisae*, we need to expend the temporal breath of observations on *R. exoculata* reproduction to better constrain its spawning seasonality and assess its degree of variability.

Although reproductive traits of vent species were suggested to be phylogenetically constrained (Tyler & Young 1999), some Alvinocaridid shrimp species also exhibit continuous reproduction. *Alvinocaris muricola* (Ramirez_Llodra & Segonzac 2006) from cold seeps off Congo is one such example, although temporally limited observations of asynchronous ovarian development could also hide seasonal patterns. In *Rimicaris chacei* from the MAR, additional observations could also challenge the current view of continuous reproduction based on time limited observations (Ramirez-Llodra et al. 2000). In contrast, *Mirocaris fortunata* exhibit many ovigerous females throughout the year, which is truly supporting continuous reproduction: Ramirez-Llodra et al. (2000) and Van Dover et al. (1996) reported many brooding females from May, June, July and September at Lucky Strike. We also repeatedly observed *M. fortunata* brooding females during multiple cruises at different periods of the year and different MAR vent fields: August 2013 and April 2015 at Lucky Strike (Momarsat cruise series), and January 2014 at Snake Pit (BICOSE cruise). To conclude, Alvinocaridid reproductive strategies are diverse making it difficult to draw general inferences for the family.

Vent species are usually believed to reproduce continuously because of the absence of photoperiodic signals at depth coupled with the high and continuous productivity at vents that would obscure variations in surface-derived food supply. Seasonal reproduction in Alvinocaridid shrimps is however not an exception. On the MAR, *Bathymodiolus azoricus* mussels spawn in January (Colaço et al. 2006, Dixon et al. 2006), and this pattern has been related to the variability in the flux of particulate food material that might trigger seasonal reproductive activity (directly in planktotrophic larvae and/or indirectly in adults). On the East Pacific Rise, vent crabs *Bythograea thermydron* brood their offsprings in spring (Perovich et al. 2003), again possibly responding to surface production cues. On the other hand, bythograeid crabs from Pacific-Antarctic Ridge vent sites were suggested to have continuous reproduction, suggesting biogeographic effects may induce variations in reproductive rhythms (Hilario et al. 2009). Similarly, the study of reproductive rhythms in other *Rimicaris* shrimps from other locations may provide insights into the mechanisms underlying spawning seasonality in *R. exoculata*.

In addition to seasonal reproduction, population size structure suggest discontinuous recruitment in *R. exoculata*, with similar patterns at both vent fields. Polymodal population structure was also noted by Gebruk et al. (1997) and Copley (1998). In addition, Shank et al. (1998) suggested discrete recruitment of juvenile cohorts based on the observation of red juveniles patches within adult swarms. Discontinuous recruitment may further suggest seasonal reproduction, although growth rates and recruitment temporality remain to be assessed before any supported conclusion can be drawn. *R chacei* in nurseries appear to be recent recruits as they are similar to those captured in nets above vents (Herring & Dixon 1998). It is less clear for *R. exoculata* juveniles observed in active emission habitats, although the abundance of lipids and isotopic signatures suggest they also have recently experienced pelagic life and were feeding (Methou et al. 2020).

### 4.4. Brooding eggs in vent fluids

The remarkable lack of *R. exoculata* brooding females in samples collected at various periods and locations, until this study, had been explained by the hypothesis of a migration of brooding females towards -yet to be found- areas with moderate environmental conditions in order to avoid exposure of their eggs to harmful vent fluids (Ramirez-Llodra et al. 2000). Such protective behavior of brooding mothers is indeed reported in many vent crustaceans. Brooding females of *B. thermydron* crabs on the East Pacific Rise were observed at the periphery of vents whereas males and non-brooding females occupied areas with high fluid venting and temperatures (Perovitch et al. 2003). In Alvinocaridid shrimps, brooding females of *A. stactophila* locate preferentially in areas with more oxygen, and less toxic sulfur compounds than those occupied by males and non-brooding females (Copley & Young 2006). In the amphipod *Ventiella sulfuris*, adults migrate at the periphery of vents to mate and brood (Sheader & Van Dover 2007). In the anomouran crab *Kiwa tyleri*, brooding females also locate at the periphery in less-constraining conditions, whereas males remain close to vent fluid exits (March et al. 2015, Thatje et al. 2015). In *R. exoculata*, however, such scenario had already been challenged by the collection of a few brooding females in dense shrimp aggregates at Logatchev (Guri et al, 2012). Our study further strengthen these observations, brooding females from TAG and Snake Pit being exclusively found in dense shrimp aggregates where temperature records were higher, and never observed (including on video footage) in cold peripheral areas where males scattered.

Females of *R. hybisae* also appear to brood their eggs directly within aggregates crawling on chimney walls with no particular protection from vent fluids. Both *R. exoculata* and *R. hybisae* are rare examples of vent species that experience vent fluid exposure during embryonic development. Indeed, many species broadcast their eggs, in which case early development occur in the water column at some distance from the most extreme part of the vent fluid mixing gradient, and brooding species usually provide protective structures for the early development of their offsprings (e.g. Reynolds et al. 2010), or relocate into milder areas.

As all developmental stages were observed in broods of *R. exoculata*, including hatching zoea (Hernández-Ávila et al. 2015), females probably remain near vent fluid exits during the whole brooding period until they release their larvae. Exposure of embryos to high temperature may accelerate their development, and shorten the brooding period, while challenging normal development due to heat stress. *In vitro* incubations of embryos removed from brooding females of *Alvinocaris longirostris* and *Shinkaicaris leurokolos*, two Alvinocaridid species inhabiting Okinawa Trough vent sites, showed increased developmental rates with increasing temperature for both species, with optimal growth at temperatures within the range experienced by their adult counterparts *in situ* (Watanabe et al. 2016). These authors showed that embryos of *S. leurokolos*, which lives near fluid exits, developed better between 10 to 20°C, and hatched within 9-12 weeks at 10°C and 3-4 weeks at 20°C. Considering the habitat of brooding females in *R. exoculata*, eggs are likely to hatch rather quickly, perhaps within a month following spawning. A relatively short brooding period in *R. exoculata* is also supported by the lack of macroscopic evidence of exceedingly high mineral deposits in the cephalothoracic cavity of ovigerous females. The short molt cycle of *R. exoculata* (10 days, Corbari et al. 2008) is believed to reset symbiotic communities of the cephalothoracic cavity sufficiently often to avoid impaired symbiosis due to mineral precipitation progressively overgrowing bacteria. In bearing-egg crustaceans, the molting cycle is interrupted during the brooding period until hatching (Correa & Thiel 2003). As females appear to spend the entire brooding period near vent fluids, a large extension of the molting cycle expose them to the risk of seeing their symbiotic bacteria drowned in mineral precipitations. However, as ovigerous females did not appear to suffer excessive mineral load of the cephalothoracic cavity, this observation also advocates for a relatively short brooding period.

Brooding within vent fluids may also sustain the bacterial community including symbiotic lineages observed on egg envelopes (Methou et al. 2019), and may thus participate in symbiont transmission to the young shrimp (Hernandez Avila et al. 2015). Another speculative hypothesis would be that mothers could imprint their offspring with vent signature by bathing their eggs within vent fluids during embryonic development, which would later help them returning to vent habitats after dispersal. Such homing process might involve the strongly developed higher brain centers of *R. exoculata* enabling the memory and navigation skills necessary to locate suitable recruitment sites (Machon et al. 2019).

### 4.5. Mating system in R. exoculata

Spatial segregation of sexes in vent species has been related to reproductive processes needing specific habitats, but it is not known whether such spatial structuration remains throughout the year or is restricted to mating and brooding periods. Segonzac et al. (1993) observed dispersed *R. exoculata* - of unknown sex - at the periphery of Snake Pit in June 1988 but more observations are needed to evaluate the temporal correlation between spatial distribution of sexes and reproductive processes. It is however unlikely that individuals resting in peripheral areas stay permanently in such habitats because of their need to supply their symbiotic bacteria with reduced compounds found in vent fluids (Ponsard et al. 2013). Moreover, the highly modified cephalothorax of *R. exoculata* adults hinder movement of the appendages, making any foraging or grazing behavior unlikely (Segonzac et al. 1993, Zbinden & Cambon 2020). Shrimps thus most likely fast when they are in IEH. Peripheral areas also harbor more predators, such as *Maractis rimicarivora* anemones (Fautin & Barber 1999, Copley et al. 2007), or fishes that are excluded from AEH, maybe because of the harsh vent conditions. The migration of *R. exoculata* males to inactive areas is therefore not likely driven by trophic needs or interspecific interactions (increased predatory risk), and given the current knowledge, the hypothesis of a reproductive behavior remains the best explanation. In addition, spermatophores at the gonopores of males were often observed in IEH, but never in males from other habitats (Pradillon, personal observation), suggesting a greater mating readiness in IEH. Mature caridean females molt just before egg extrusion (Bauer 2004), and vent shrimps may migrate briefly towards a milder habitat at this stage of increased vulnerability. Males in IEH would thus have better chances to encounter females just after their prespawning molt and mate successfully. After mating and egg extrusion, females would return to AEH and brood their eggs, while the fate of males remains uncertain. Males will however probably have to return to AEH at some point to fulfill their nutrition needs.

Body sizes and male weaponery in crustaceans have been associated with their mating system (Correa & Thiel 2003, Baeza & Thiel 2007, Asakura 2009). In free-living caridean shrimps, larger males are usually observed in mating systems that involve sexual competition for females or precopulatory mate guarding (Bauer 1996, Correa & Thiel 2003, Asakura 2009). In contrast, the lack of sexual competition in “pure searching” mating systems or long-term mate (monogamy or semi-monogamy) is generally associated with similar or smaller-sized males (Correa & Thiel 2003, Bauer 1996). Although *R. exoculata* males were on average larger than females, this was due to the large proportions of small females. Considering only sexually mature females, i.e. those females actually involved in the courtship and mating processes, then sex size differences do no longer exist, and females are even slightly larger than males. In other vent carideans such as *Lebbeus virentova,* females tend to be larger than males (Nye & Copley 2014), and a similar trend has been proposed for the cold-seep alvinocaridid *A. muricola*, based on maximum sizes of males and females (Ramirez-Llodra & Segonzac 2006). In *R. hybisae,* males and females exhibit similar sizes (Nye et al. 2013). In addition, in *R. exoculata* there is a lack of sexual dimorphism in secondary characters associated with male competition (e.g. increase in cheliped size and cephalotorax). We thus hypothesize that a pure searching model could better describe the mating system in *R. exoculata*. In this model, males search for receptive females just after their reproductive molt using mostly tactile signals or pheromones (Bauer 1996, 1976). However, looking for tactile and chemical cues within a dense aggregation of congeners close to vent emissions does not seem optimal. A migration to the vent periphery to courtship and mate could increase the chance of male-female recognition and ensure mating success. In his description of the mating behavior of the Hippolytidae *Heptocarpus pictus*, a caridean species exhibiting a pure searching mating model, Bauer (1976) described a suite of events including one called “straddle” when the male clutches the female with his walking legs. We observed *R. exoculata* pairs with similar behavior several times *in situ* (in all habitats, but mainly at the periphery of AEH or in DEH) (Supplementary video 1), which further supports the pure searching mating model in this species.

### 4.6. A proposed life cycle scenario for Rimicaris exoculata

Our examination of population structure and reproductive features across different habitats allows us to draw a hypothetical scenario for the benthic phase of the life cycle of *R. exoculata* (Fig. 6). In our observations, juveniles of the species appear to recruit directly in dense aggregations crawling on vent active structures. Visually, they can be distinguished from adults and subadults by their bright red color. Although in this study “nurseries” near diffusion areas belonged to *R. chacei,* juveniles of *R. exoculata* were observed forming gatherings within dense assemblages (Pradillon, personal observation and Shank et al. 1998), perhaps related to small-scale environmental conditions within AEH. Considering all observations gathered so far in this study and others (Shank et al. 1998, Gebruck et al. 2000, Methou et al. 2020), juveniles found in AEH are the smallest observed and identified with confidence for *R. exoculata*. We cannot exclude that we are still lacking an earlier benthic stage, as smaller alvinocaridid juveniles - without species assignment – were caught in bottom trawls (Herring & Dixon 1998). However, isotopic signatures of the smaller benthic stages of *R. exoculata* reported by Methou et al. (2020) are reflecting a pelagic lifestyle and advocate for relatively recent recruitment of those stages.

**Figure 6.**
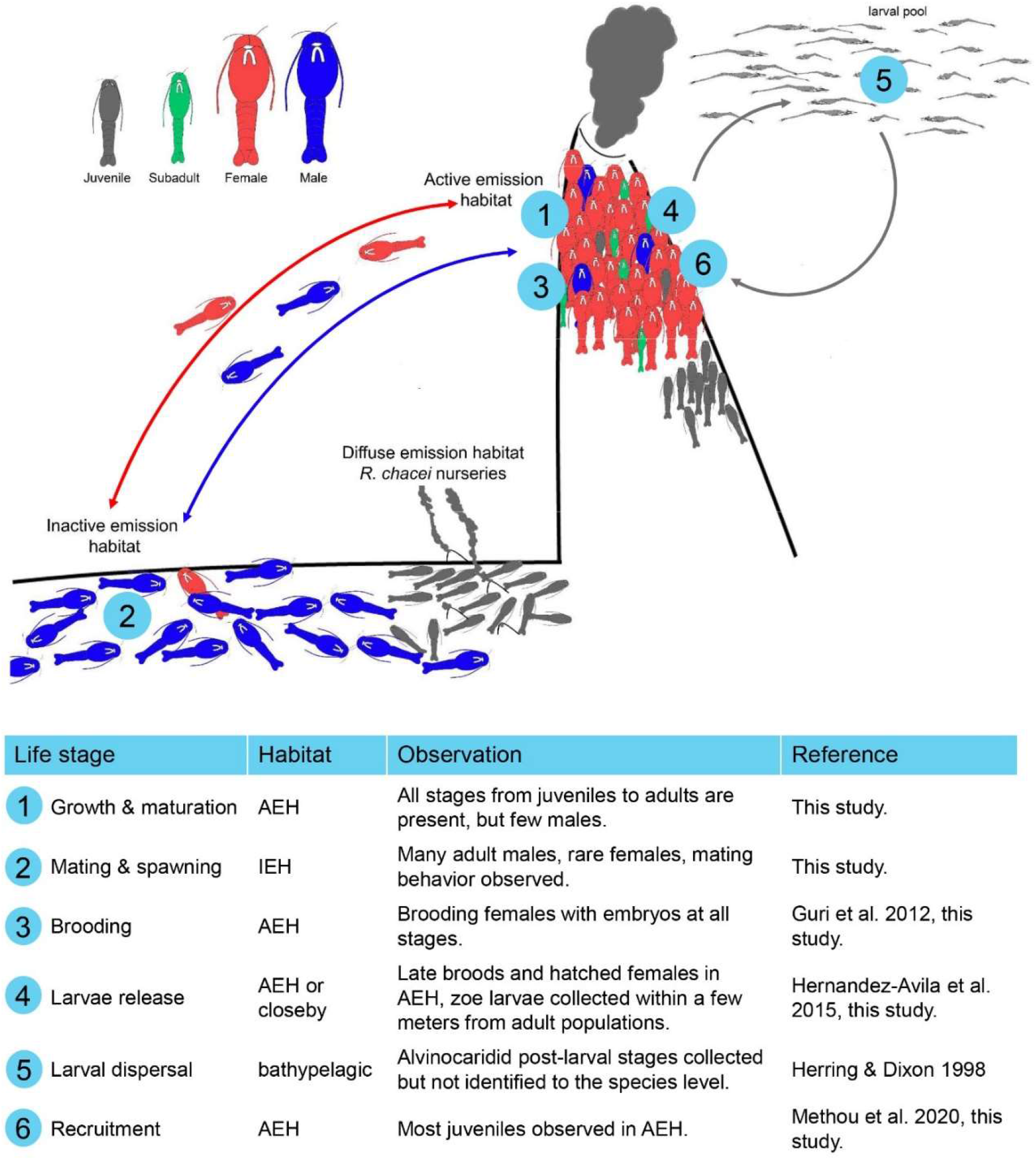
Proposed model of habitat use through the life cycle of the vent alvinocaridid shrimp *Rimicaris exoculata*

After recruitment, juveniles shift towards a chemosynthetic nutrition, and grow to the subadult and adult stages. While small adults appear to stay within AEH as they grow, mature and reach reproduction size, sexually mature reproductive adults could migrate in less active parts of vent fields for mating. Sexually mature males would move there, reaching a position where they may more “easily” find sexually receptive females that move out of dense aggregations to molt, mate and extrude their eggs. No competitive or guarding behavior was observed in males, and their size broadly similar to the size of females, as well as their lack of weaponery, do not argue for competition. Instead, males appear rather inactive, and exhibit behavior resembling the “pure searching” model where males contact females to find and mate with receptive females. Brooding would occur entirely within vent mixing gradients and last for a few weeks before release of zoea larvae. These larvae would then disperse and mix within bathypelagic waters, developing and feeding for a while on pelagic food items until they reach a large post-larval stage before returning to a benthic and chemosynthetic life style at vents.

## Credit author statement

Iván Hernández-Ávila : Conceptualization, methodology, formal analysis, investigation, writing – original draft.

Marie-Anne Cambon-Bonavita : Resources, supervision, writing – review & editing.

Jozée Sarrazin : Resources, writing – review & editing.

Florence Pradillon : Conceptualization, methodology, resource, supervision, writing – review & editing.

## Acknowledgment

This research was supported by Ifremer REMIMA program, EU Seventh Program for Research, Technological Development and Demonstration Activities, MIDAS grant 603418, Fundación Gran Mariscal de Ayacucho PhD grant E-223-85-2012-2 to IHA and Campus France grant 796045K to IHA. We also thank the captain and crew of the oceanographic cruise BICOSE-2014 (DOI: 10.17600/14000100), as well as the pilots of the ROV Victor 6000. We also thank M. Segonzac for his help on juvenile sorting during the BICOSE cruise.

