## supplemental material for "Population structure and reproduction of the alvinocaridid shrimp *Rimicaris exoculata* on the Mid-Atlantic Ridge: variations between habitats and vent fields"

**Supplementary Table 1**. Individuals used for barcoding.

| Individual label | Taxa/Life stage | Vent field | Sample | Accession number |
|---|---|---|---|---|
| CHj4/J9/J12/J68/Rej3 | Rimicaris juvenile | TAG | S13 | KU948492 |
| CHj5/J7/J8/J11/J16/J20/J66/J67/ J69/J73/Rej1/Rej2/Rej6/Rej7 | Rimicaris juvenile | TAG | S13 | KU948491 |
| J1 | Rimicaris juvenile | TAG | S13 | KU948497 |
| J4/J15 | Rimicaris juvenile | TAG | S13 | KU948500 |
| J5/J6 | Rimicaris juvenile | TAG | S13 | KU948498 |
| J10 | Rimicaris juvenile | TAG | S13 | MW052711 |
| J13 | Rimicaris juvenile | TAG | S13 | MW052712 |
| J18/J77 | Rimicaris juvenile | TAG | S13 | KU948499 |
| J70 | Rimicaris juvenile | TAG | S13 | KU948494 |
| J71 | Rimicaris juvenile | TAG | S13 | MW052713 |
| J72 | Rimicaris juvenile | TAG | S13 | KU948491 |
| J74 | Rimicaris juvenile | TAG | S13 | KU948506 |
| J75 | Rimicaris juvenile | TAG | S13 | KU948507 |
| J76 | Rimicaris juvenile | TAG | S13 | KU948493 |
| J103 | Rimicaris juvenile | TAG | S13 | KU948508 |
| J104 | Rimicaris juvenile | TAG | S13 | KU948503 |
| J21/J22/J23/J28/J30/J34/J35/ J37/J39/J40/J79/J83/J85/J86/ J87/Rej9/Rej10 | Rimicaris juvenile | TAG | S14 | KU948491 |
| J24/J27/J29/J31/J32/J38/J81/ J84/Rej8 | Rimicaris juvenile | TAG | S14 | KU948492 |
| J25 | Rimicaris juvenile | TAG | S14 | KU948504 |
| J26 | Rimicaris juvenile | TAG | S14 | KU948507 |
| J33 | Rimicaris juvenile | TAG | S14 | KU948508 |
| J36 | Rimicaris juvenile | TAG | S14 | KU948496 |
| J80 | Rimicaris juvenile | TAG | S14 | KU948502 |
| J89 | Rimicaris juvenile | TAG | S14 | KU948495 |
| J109 | Rimicaris juvenile | TAG | S14 | KU948505 |
| Rej11 | Rimicaris juvenile | TAG | S14 | KU948501 |
| Rej12 | Rimicaris juvenile | TAG | S14 | KU948499 |
| J88 | Rimicaris juvenile | TAG | S14 | MW052714 |
| Rej5 | Rimicaris juvenile | TAG | S7 | MW052724 |
| J61 | Rimicaris juvenile | TAG | S7 | MW052716 |
| J62 | Rimicaris juvenile | Snake Pit | S6 | MW052717 |
| J64 | Rimicaris juvenile | TAG | S7 | MW052723 |
| J65 | Rimicaris juvenile | TAG | S7 | MW052719 |
| Ra1 | *R. exoculata* adult | TAG | S9 | KY632681 |
| Ra2 | *R. exoculata* adult | TAG | S8 | KY632682 |
| Cha1 | *R. chacei* adult | Snake Pit | PL05-Aspi4 | KY632687 |
| Cha2 | *R. chacei* adult | Snake Pit | PL05-Aspi4 | KY632688 |
| C3097 | copepod | TAG | S8 | MW065791 |

**Supplementary Table 2**. Carapace length (mean and range, mm) of *R. exoculata* specimens collected at different vent fields and habitats. J: juveniles; <OSD: subadults; F: females (non-brooding); OF: ovigerous females; M: males. ND: no data (damaged specimen).

| Habitat | Vent field | Sample | J | <OSD | F | OF | M | Total per Habitat/Vent field |
|---|---|---|---|---|---|---|---|---|
| **Active Emission** | | | | | | | | |
| | Snake Pit | S01 | 9 | 9.01 (7.8-9.5) | 12.68 (10-19.4) | 17 | 14.04 (12-16.3) | |
| | Snake Pit | S02 | 8.71 (7.8-10.1) | 8.98 (7-9.9) | 11.19 (10-20.3) | 17.50 (17.3-17.7) | 13.50 (10.5-16.6) | |
| | Snake Pit | S03 | 8.54 (7.5-9.4) | 9.16 (7.5-9.9) | 11.89 (10-20.4) | 17.05 (15-19.9) | 14.06 (9.9-17.1) | |
| | Snake Pit | S04 | 8.34 (7.3-9.3) | 8.88 (7.2-9.9) | 11.14 (10-14.7) | | 12.47 (10-14.7) | |
| | Snake Pit | S05 | 8.22 (5.6-9.2) | 8.60 (7.5-9.7) | 12.93 (10-24.4) | 16.2 | 15.29 (10-19.1) | |
| | Snake Pit | S06 | 8.94 (8.2-10.3) | 9.01 (7.1-9.9) | 13.30 (10-20.6) | 17.83 (15.5-20.5) | 14.86 (10.6-18.2) | 11.12 (5.6-24.4) |
| | TAG | S07 | 9.18 (8.2-10.3) | 9.05 (7-9.9) | 12.35 (10-18.6) | 15.52 (12.1-17.1) | 14.38 (11.9-17.2) | |
| | TAG | S08 | 9.7 | 9.47 (8.8-9.9) | 13.23 (10-20.2) | 15.03 (12-17.5) | 15.40 (13.5-18.6) | |
| | TAG | S09 | 9.17 (8.7-9.5) | 9.30 (7.5-9.8) | 14.65 (10-19.5) | 17.65 (14.3-20.6) | 16.62 (11.4-18.4) | 12.57 (7-20.6) |
| **Inactive Emission** | | | | | | | | |
| | TAG | S10 | 9.1 | | 11.25 (10.8-11.7) | | 15.51 (12.2-18.9) | |
| | TAG | S11 | | 9.55 (9.2-9.9) | 13.75 (11.3-16.2) | ND | 15.78 (12.6-17.5) | |
| | TAG | S12 | | | 15.2 | | 15.98 (12.9-18) | 15.45 (9.1-18.9) |
| **Diffuse Emission** | | | | | | | | |
| | TAG | S13 | 5.18 (4.4-6.5) | | | | | |
| | TAG | S14 | 5.08 (4.5-5.8) | | | | | 5.12 (4.4-6.5) |

**Supplementary Table 3**: ANOVA of body sizes of adult stages between sex and vent fields in the active emission habitat (AEH). In bold, significant results.

| Source | df | SS | MS | F | p |
|---|---|---|---|---|---|
| Vent field | 1 | 1.001 | 1.001 | 4.095 | 0.083 |
| **Sex** | **1** | **1.448** | **1.448** | **80.712** | **<0.001** |
| **Samples (Vent)** | **7** | **2.166** | **0.309** | **15.764** | **<0.001** |
| Vent x Sex | 1 | 0.002 | 0.002 | 0.088 | 0.772 |
| Sample (Vent) x Sex | 7 | 0.122 | 0.017 | 0.889 | 0.514 |
| Residual | 2103 | 41.281 | 0.02 | | |
| Total | 2120 | 55.855 | | | |

**Supplementary Table 4**. ANOVA of male body sizes between the different habitats at the TAG vent field. In bold, significant results.

| Source | df | SS | MS | F | p |
|---|---|---|---|---|---|
| Habitat | 1 | 0.010 | 0.010 | 0.163 | 0.707 |
| **Samples (ha)** | **4** | **0.332** | **0.080** | **10.856** | **<0.001** |
| Residual | 156 | 1.157 | 0.007 | | |
| Total | 161 | 1.478 | | | |

**Supplementary Table 5**. ANOVA of embryo volume associated between vent fields, embryo stages and parental females. In bold, significant results.

| Source | df | SS | MS | F | p |
|---|---|---|---|---|---|
| **Vent** | **1** | **0.06043** | **0.06043** | **10.715** | **0.002** |
| **Stage** | **2** | **0.24414** | **0.12207** | **21.648** | **<0.001** |
| V x St | 2 | 0.01511 | 0.00755 | 1.340 | 0.267 |
| **Female (V x St)** | **94** | **0.52826** | **0.00562** | **15.319** | **<0.001** |
| Residual | 896 | 0.32869 | 0.00037 | | |
| Total | 995 | 1.18940 | | | |

**Supplementary Figure 1**. Shrimp assemblages sampled in AEH at Snake Pit (A: S1, B: S2 & S3, C: S4, D:  S5, E: S6), and TAG (F: S7, G: S8, H: S9). (Ifremer, ROV Victor6000, BICOSE 2014)

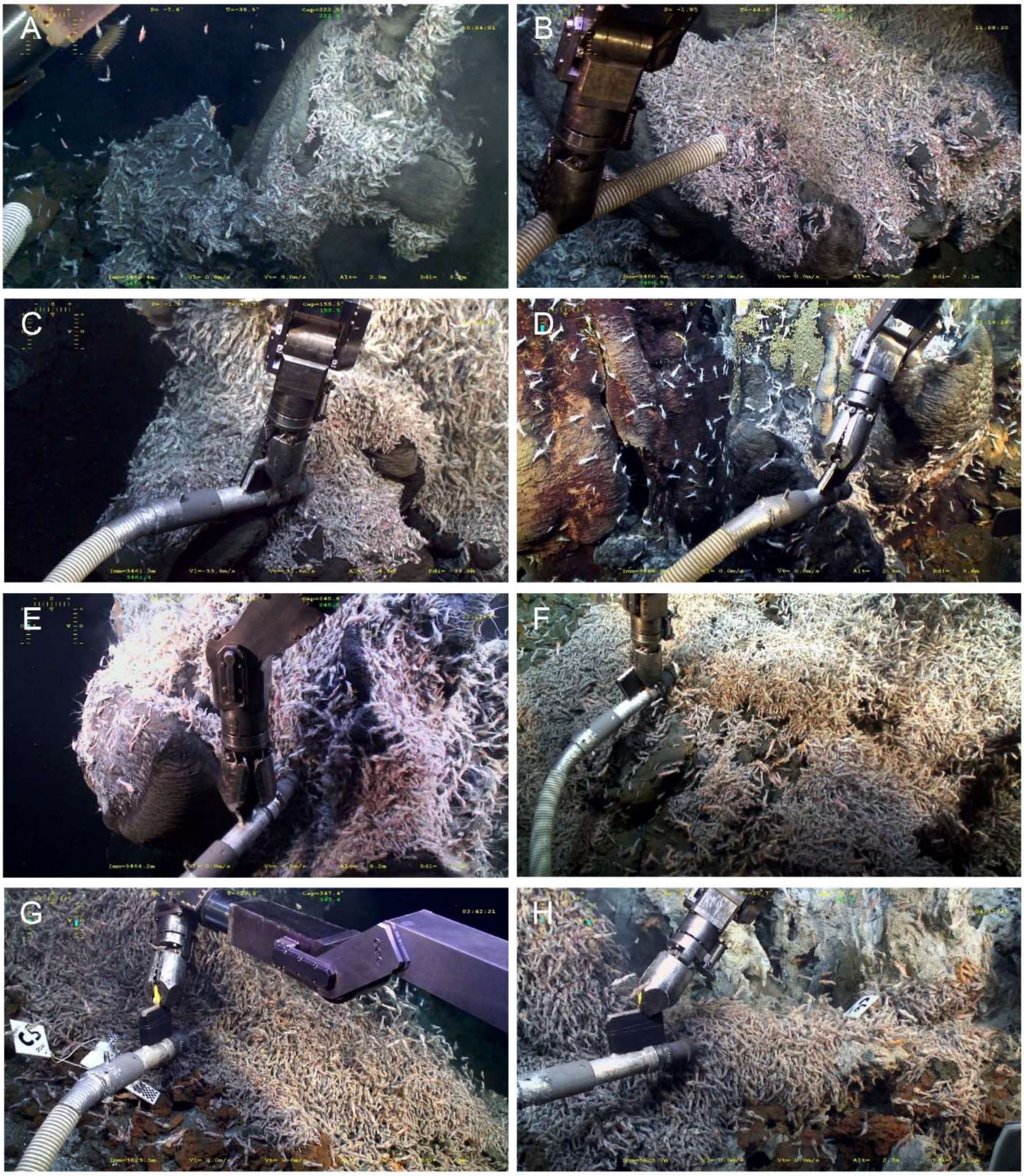

**Supplementary figure 2**. Shrimp assemblages sampled in the IEH at TAG (A: S10, B: S11, C: S12), and in the DEH at TAG (D: S13, E: S14) and Snake Pit -Moose edifice (F: sampling of early juveniles -yellow arrow- among *Bathymodiolus puteoserpentis* mussels). (Ifremer, ROV Victor6000 BICOSE 2014)

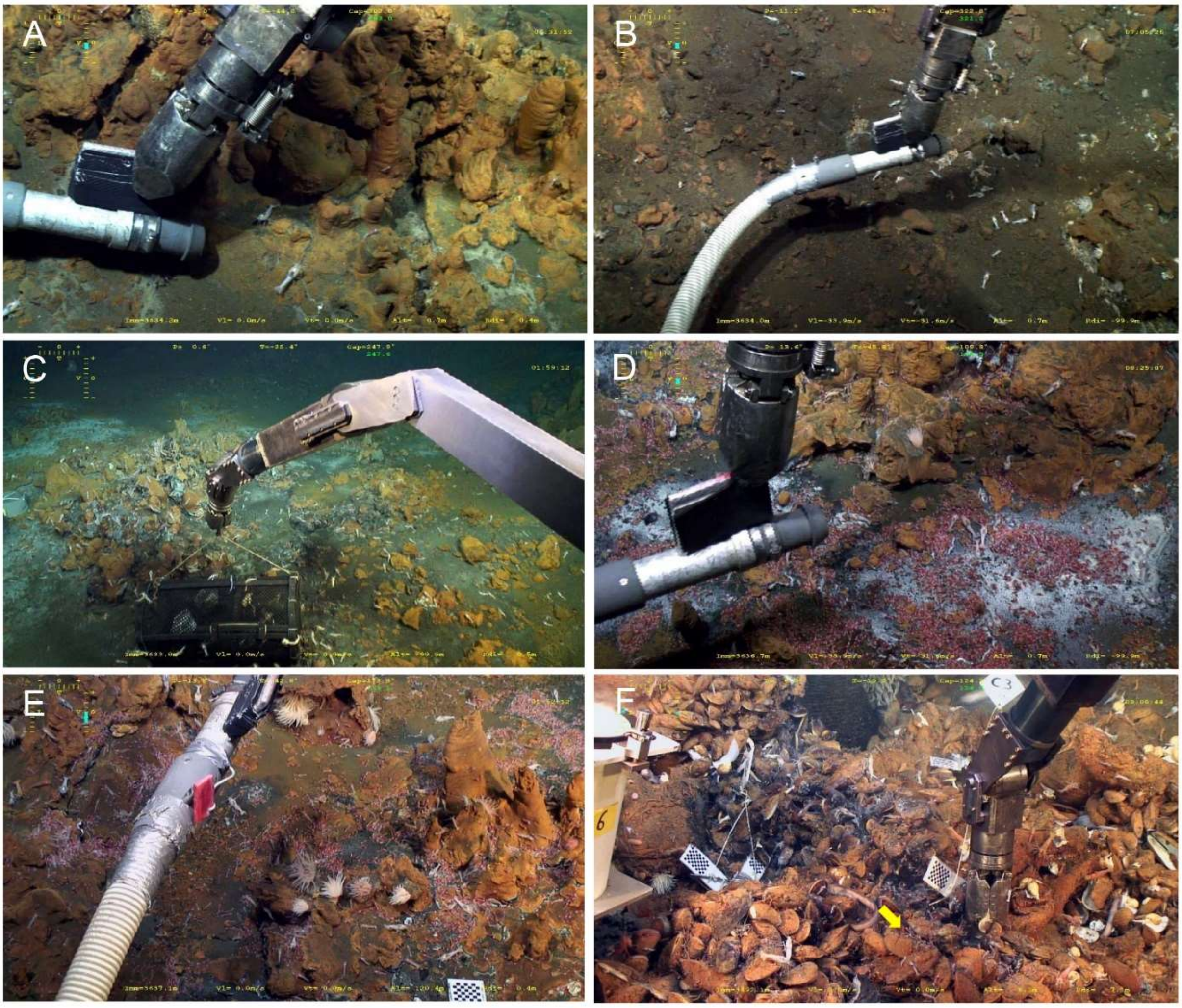

**Supplementary figure 3**. Onset of sexual differentiation (OSD) determination (sexual determination: ratio of individuals with sufficient gonad development to allow sex identification).

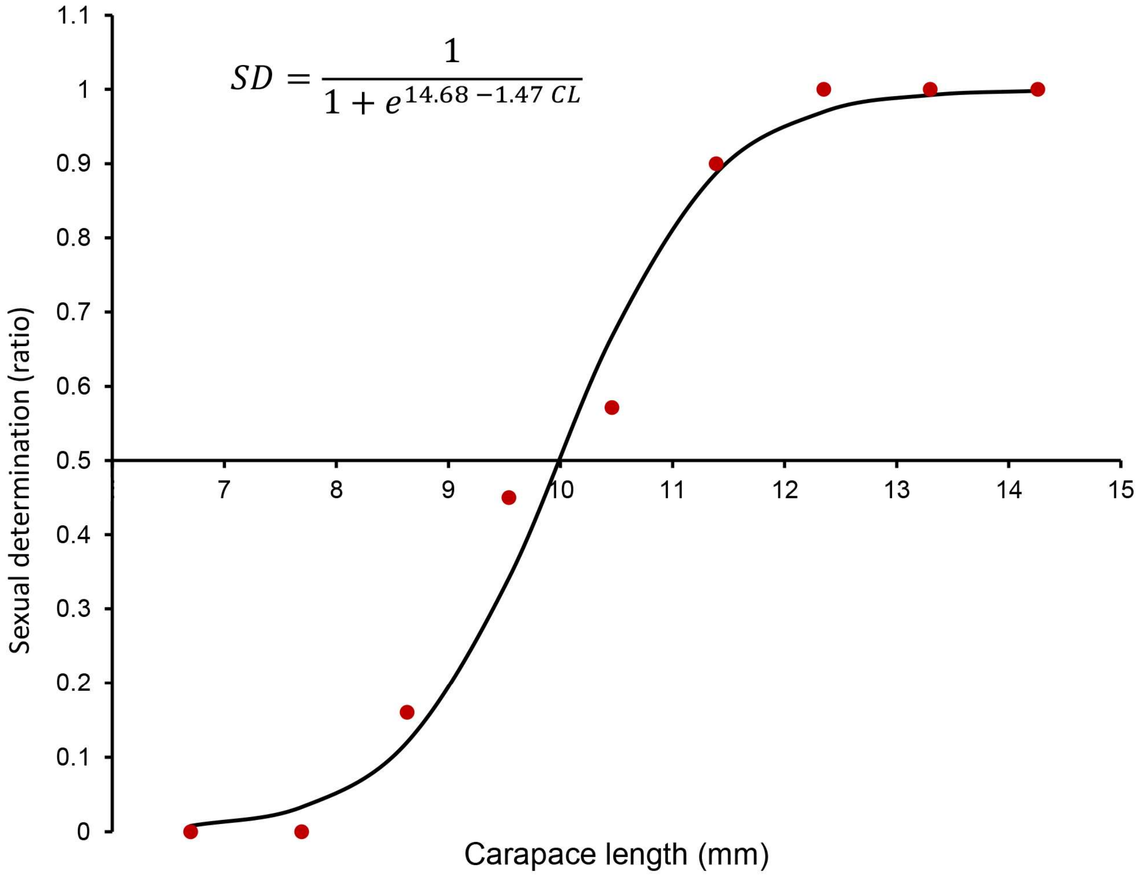

**Supplementary figure 4**. Onset of effective sexual maturity (ESM) determination. Effective sexual maturity ratio = proportion of ovigerous females per size class corrected by the maximum proportion of ovigerous females per size class.

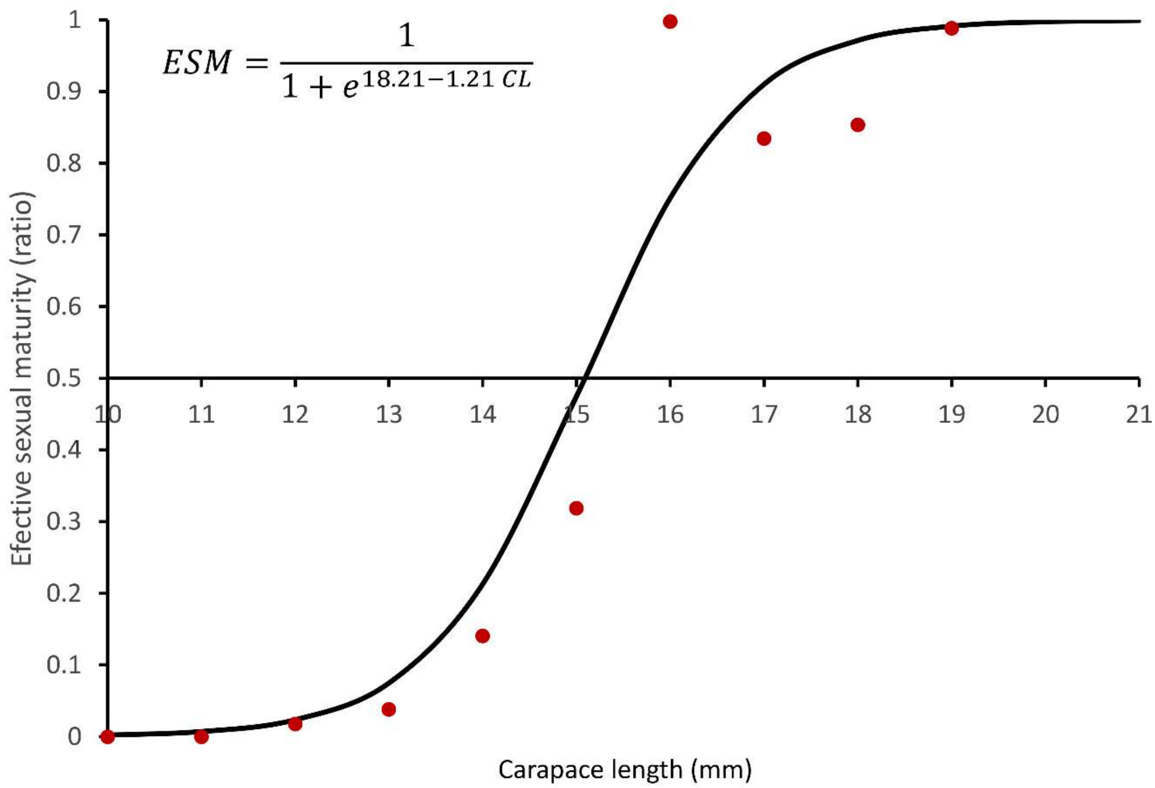

**Supplementary figure 5**. Size distribution of *Rimicaris exoculata* shrimps from the different habitats at TAG.

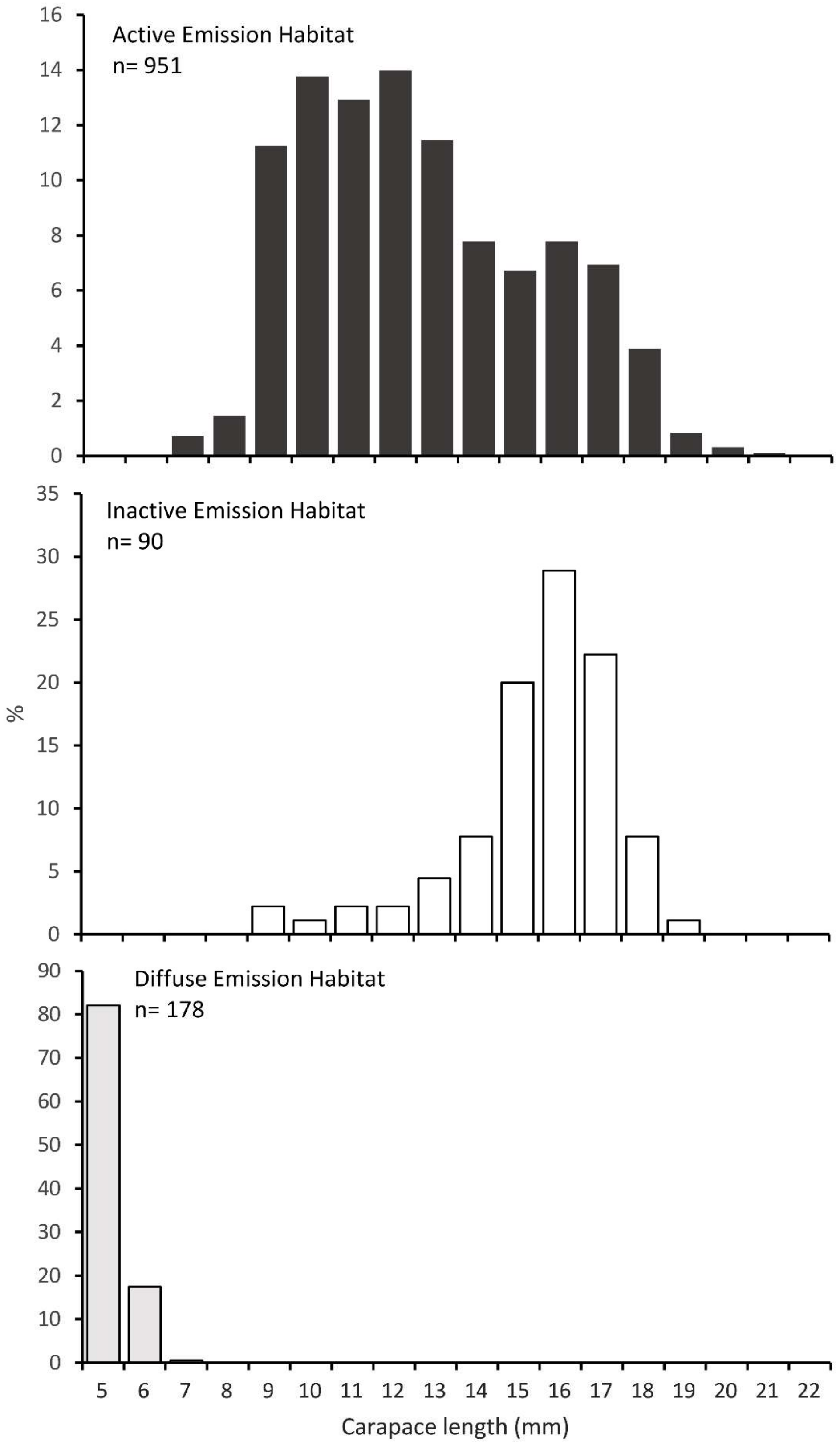

**Supplementary figure 6**. Modal decomposition of *Rimicaris exoculata* populations in AEH.

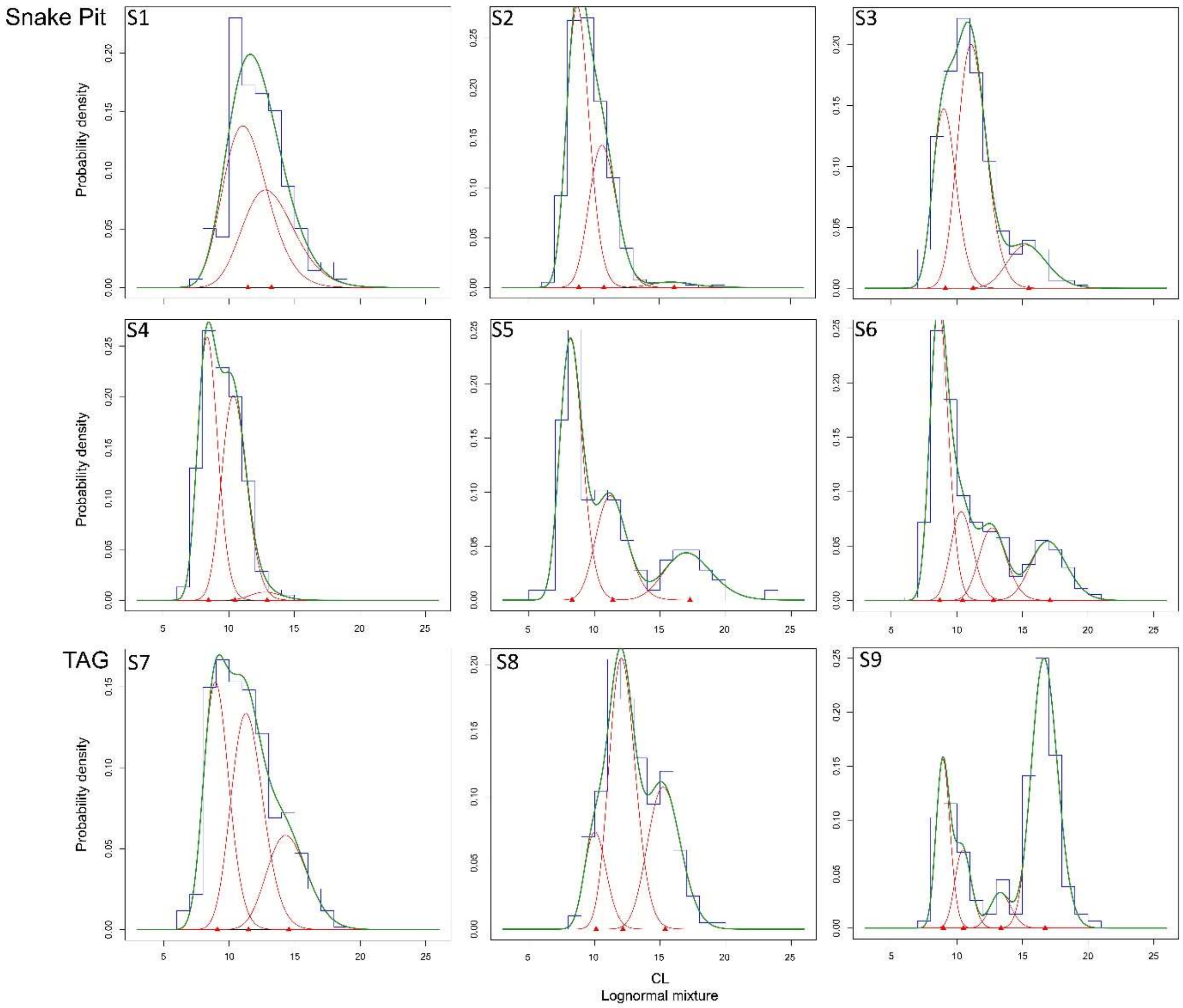

**Supplementary figure 7**. Neighbour-Joining tree of *Rimicaris* juvenile COI barcodes (443 bp). The blue clade gathers sequences assigned to *R. chacei*, and the yellow clade gathers sequences assigned to *R. exoculata*. 72 specimens collected in juvenile aggregations (nurseries) in diffuse emission habitat (DEH) and 5 juveniles selected from populations sampled on active emission habitats (AEH) are included. Adults of *R. exoculata* (Rex) and *R. chacei* (Rcha) were also sequenced for reference and marked with black dots. Accession numbers are in Supplementary Table 1.

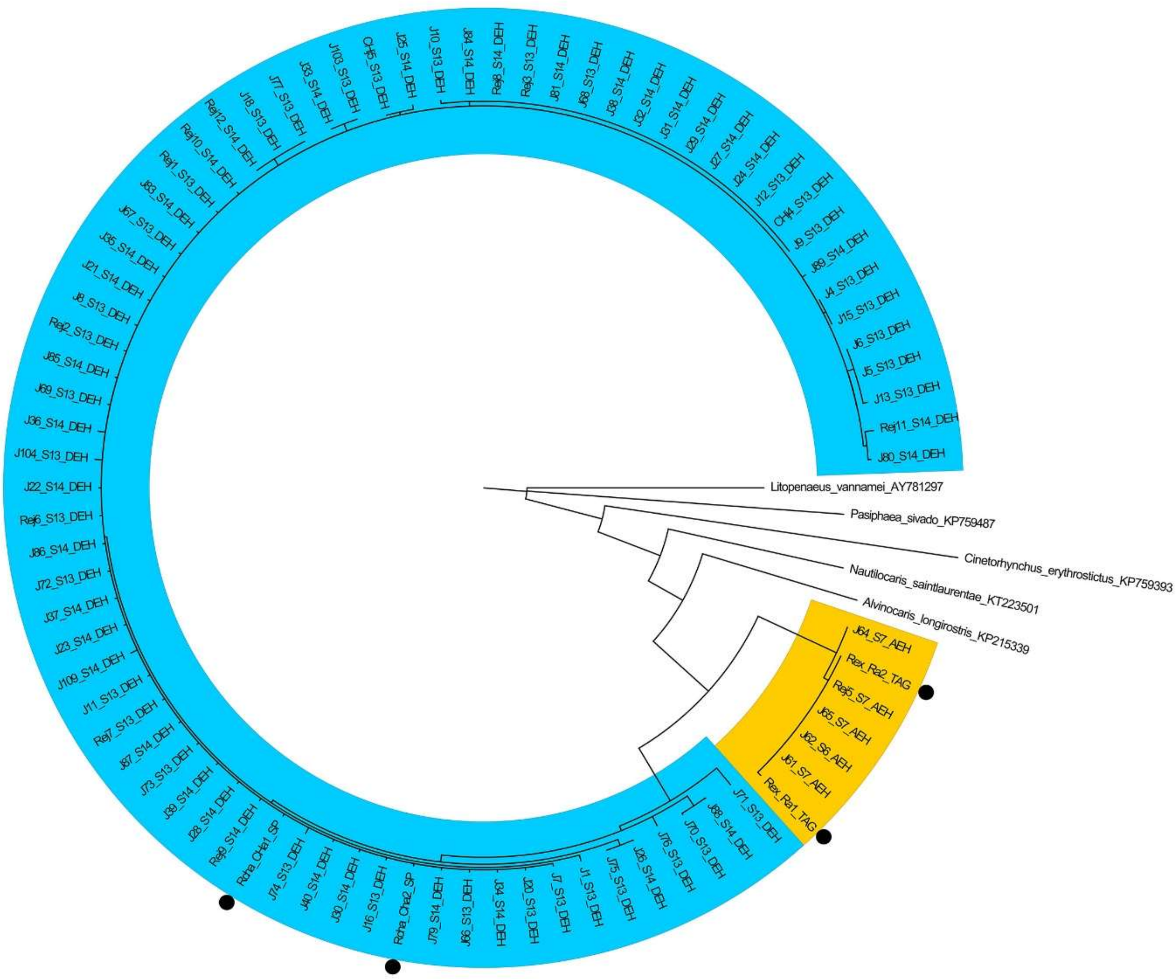

**Supplementary figure 8**. Developmental stages distribution in brooding females of each AEH sample. Samples with less than 2 brooding females are not represented.

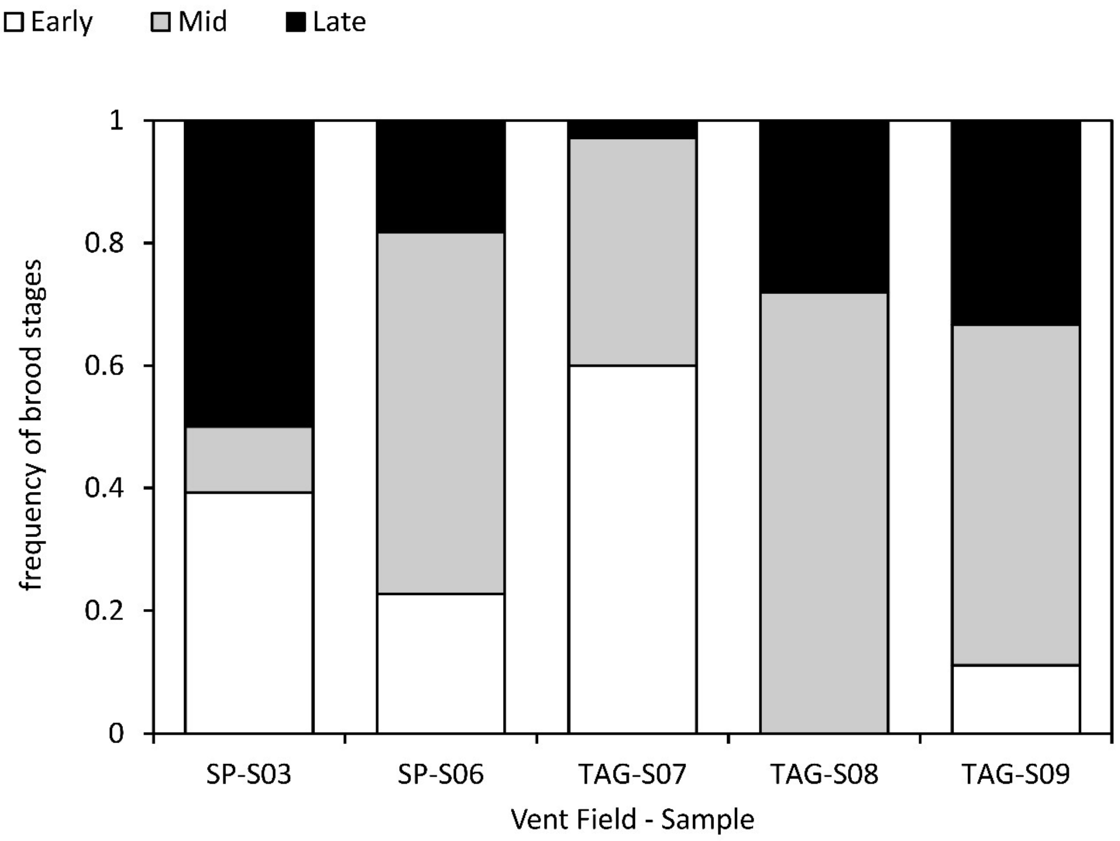


**Supplementary figure 9**. Neighbour-Joining tree of copepod COI barcodes.

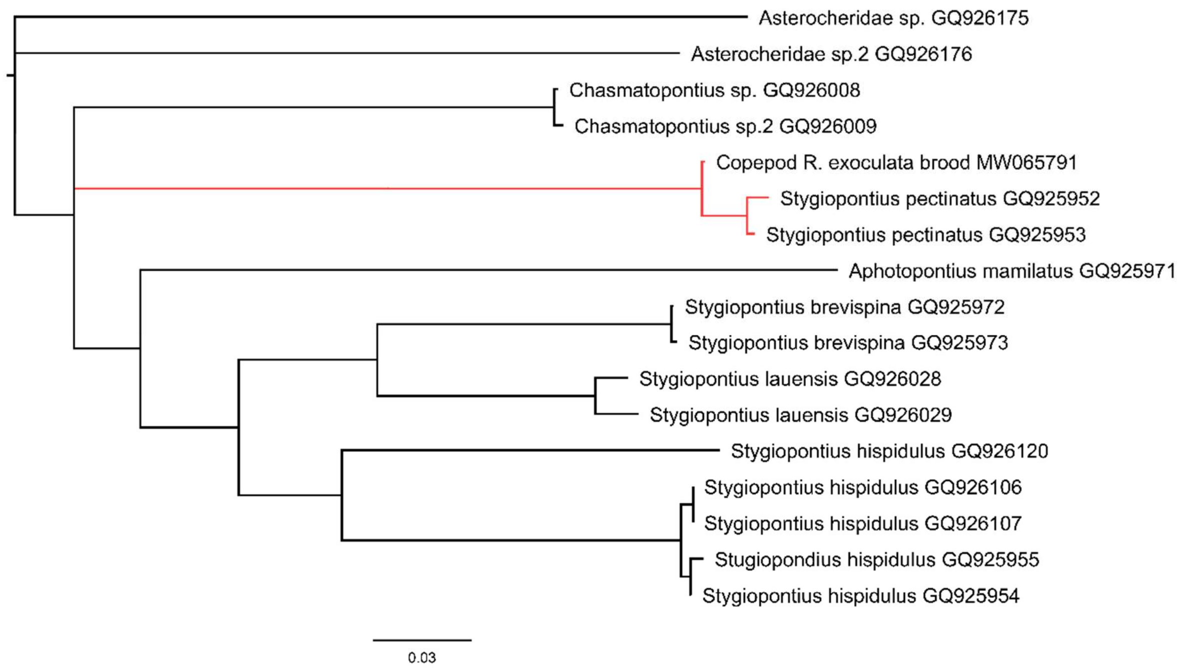

**Supplementary Figure 10**. Proportion of brooding females with broods infested with *Stygiopontius pectinatus* along embryonic development. Data from both vent fields were pooled.

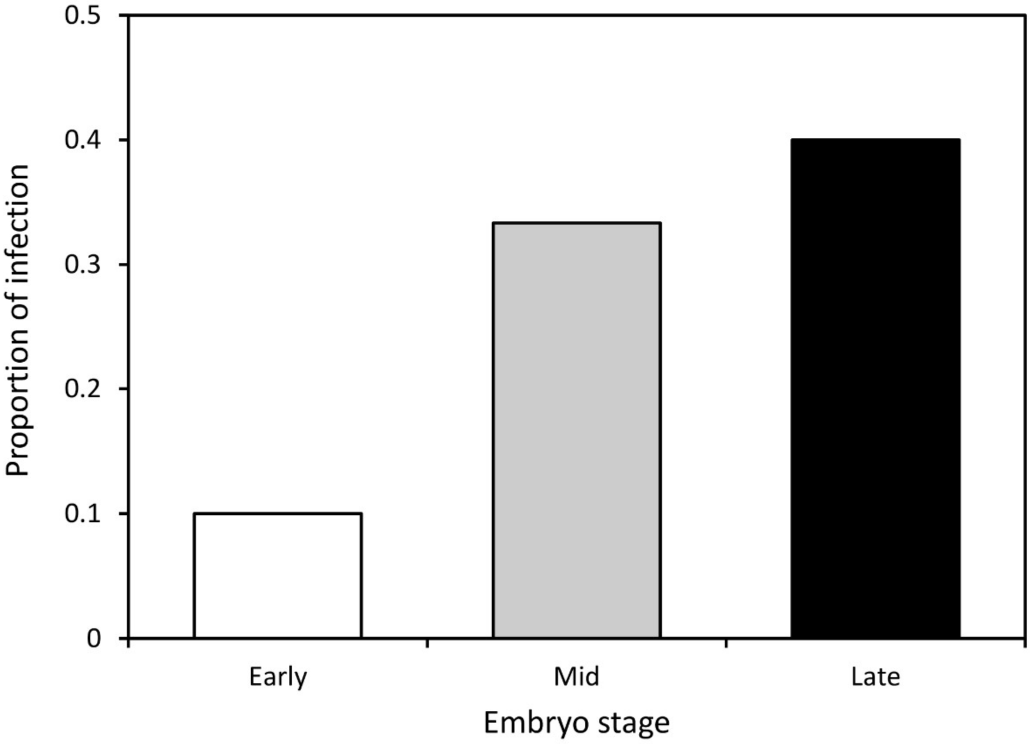


**Supplementary video 1.** ROV VICTOR6000 video acquisitions in diffuse emission habitat or at the edge of active emission habitat showing putative mating behaviour in *R. exoculata*. The images show couples of adult shrimps one over the other, resembling a straddle position (putative male on top of the putative female) described for shrimps with pure searching mating system (Bauer 1976). 0’00- 0’35 : DEH at TAG, a couple of *R. exoculata* in the center of the image crawling among *R. chacei* juveniles, as well as other adults of *R. exoculata* or *R. chacei*. 0’36-0’54 : near AEH at TAG, while the submersible is holding a temperature probe within the aggregate, a couple forms on the claw of the submersible, including a mature female (bright pink gonads visible dorsally). 0’55-1’06 : near AEH at Snake Pit: a couple forms, including a mature female (bright pink gonads) on the right of the image on bare substratum.
